# Cerebral ischaemic stroke results in altered mucosal IgA responses and host-commensal microbiota interactions

**DOI:** 10.1101/2024.06.23.600084

**Authors:** Madeleine Hurry, Raymond Wong, Laura Díaz-Marugan, Bianca De Leon, Corinne Benakis, Ari Waisman, Stuart M. Allan, Catherine B. Lawrence, David Brough, Matthew R. Hepworth

**Affiliations:** School of Biological Sciences, Faculty of Biology, Medicine and Health, Manchester Academic Health Science Centre, University of Manchester, M13 9PL, United Kingdom; Lydia Becker Institute of Immunology and Inflammation, University of Manchester, United Kingdom; Geoffrey Jefferson Brain Research Centre, University of Manchester, Manchester, UK; Institute for Stroke and Dementia Research (ISD), University Hospital, LMU Munich, Munich, Germany; Institute for Molecular Medicine, University Medical Center of the Johannes Gutenberg-University Mainz, Mainz, Germany

**Keywords:** Stroke, MCAO, Commensal Microbiota, Immunoglobulin A, intestine, immune

## Abstract

Stroke is a devastating neurological event with a high risk of mortality, but also results in long-term sequalae in survivors that extend beyond the central nervous system. Notably, these include gastrointestinal dysfunction and alterations in the commensal microbiota in both patients and mouse models, which have been suggested to contribute to secondary infection and poor outcome following stroke. Strikingly changes in commensal microbial community composition occur rapidly in both humans and animal models following stroke and correlate with disease severity. Despite these observations the underpinning mechanisms that drive alterations in the microbiota post-stroke remain poorly understood. The gastrointestinal tract is home to a complex network of tissue-resident immune cells that act constitutively to maintain microbial community and prevent bacterial-driven inflammation. Here we demonstrate that mice subjected to ischaemic stroke exhibit alterations in the intestinal immune system, most notably in antibody secreting B cells and the production of Immunoglobulin A (IgA) – a major effector response against commensal microbes. Mice lacking secretory IgA binding to commensal bacteria exhibit a partial reversion of stroke-induced changes in microbiota composition. Notably we also report increases in B cell and IgA-producing plasma cell frequencies in the brain and meninges following stroke. Together these findings demonstrate stroke is associated with perturbations in antibody producing immune responses both in mucosal tissues and the CNS following stroke, which in part explain stroke-induced changes in the intestinal microbiota. A mechanistic understanding of the immunological basis of stroke-associated pathologies in the periphery may open new avenues to manage the secondary complications and long-term prognosis of patients suffering from neurological disease.

## Introduction

Stroke is a devastating neurological event that is the second leading cause of death worldwide, and a significant cause of morbidity and disability (Feigin et al., 2022). Whether ischaemic or haemorrhagic in origin, stroke leads to a plethora of primary and secondary pathologies, including motor and sensory deficits, cognitive impairment, behavioural change and mood disorders (Stroke Association, 2018; Feigin et al., 2022; Feigin et al., 2014; Iadecola and Anrather, 2011; Iadecola et al., 2020). Additionally, up to one third of patients will experience post-stroke infection, most commonly pneumonia or urinary tract infection, adversely affecting survival and recovery (Westendorp et al., 2011). Moreover, up to 50% of stroke patients also experience gastrointestinal complications, including dysphagia, constipation, gastrointestinal bleeding, incontinence, and incomplete bowel emptying (Benakis and Liesz, 2022; Camara-Lemarroy et al., 2014; Lin et al., 2013; Schaller et al., 2006).

The gastrointestinal tract is host to a complex community of commensal microbes that have critical roles in mammalian health and disease (Lynch and Pedersen, 2016). Increasing evidence suggests that stroke also results in rapid changes in the composition of the gastrointestinal microbiota in both mice and humans (Houlden et al., 2016; Singh et al., 2016; Stanley et al., 2018), which is associated with increased mortality and worsened neurological function (Sun et al., 2021; Xia et al., 2019; Xu et al., 2019). In experimental models of stroke in rodents, changes to the commensal microbiota are also associated with alterations in intestinal motility and loss of barrier integrity (Benakis and Liesz, 2022; Benakis et al., 2020a; Brichacek et al., 2020; Crapser et al., 2016; Delgado Jimenez and Benakis, 2021; Diaz-Marugan et al., 2023; Durgan et al., 2019; Houlden et al., 2016; Singh et al., 2016; Stanley et al., 2016; Stanley et al., 2018).

Despite these observations, the underlying factors that drive loss of intestinal function and modulation of the microbiota following stroke remain poorly understood. Within the gastrointestinal tract a layered network of tissue-resident immune cells act constitutively to reinforce the intestinal barrier and to regulate host interactions with the intestinal microbiota (Belkaid and Hand, 2014). Increasing evidence suggests stroke is associated with systemic immune dysregulation mediated in part by aberrant sympathetic nervous signalling (Crapser et al., 2016; Houlden et al., 2016; Schulte-Herbruggen et al., 2009; Stanley et al., 2016; Tuz et al., 2022). However, the extent to which stroke causes altered mucosal immune function - and the degree to which loss of homeostatic immune control impacts upon the host microbiota – remains unclear.

One major mechanism through which the intestinal immune system regulates commensal microbial composition is via the production of Immunoglobulin A (IgA) (Bunker and Bendelac, 2018; Huus et al., 2021; Macpherson et al., 2008; Pabst, 2012; Pabst and Slack, 2020; Rollenske et al., 2021). IgA regulation of the microbiota is critical for mammalian health, while alterations in intestinal IgA responses can drive intestinal inflammation (Hansen et al., 2019), increased susceptibility to infection (Pabst and Slack, 2020), and metabolic dysfunction (Luck et al., 2019). Critically, IgA deficiency has been shown to result in altered gut microbiota composition in both mouse models and humans (Catanzaro et al., 2019; Rigoni et al., 2016; Suzuki et al., 2004). Here we describe acute changes in the intestinal IgA response are associated with stroke in mice and propose that changes in intestinal immune responses may in part explain the mechanistic basis of stroke-induced dysbiosis.

## Results

### Stroke induces changes in microbiota composition and the intestinal transcriptome

To begin to investigate whether stroke is associated with changes in intestinal immune responses and perturbation of the commensal microbiota we subjected mice to Middle Cerebral Artery Occlusion (MCAO) (Figure 1A). In line with prior findings (Crapser et al., 2016; Stanley et al., 2016), we confirmed rapid loss of intestinal barrier integrity and observed marked changes in commensal microbial composition, including a trend towards reduced microbial diversity (Shannon Index) and a significant decrease in *Firmicutes* to *Bacteroides* ratio as early as 24-hours following stroke (MCAO) – which was not observed in animals undergoing sham surgery, or in naïve mice (Figure 1B-E, Fig. S1A-B).

**Figure 1.**
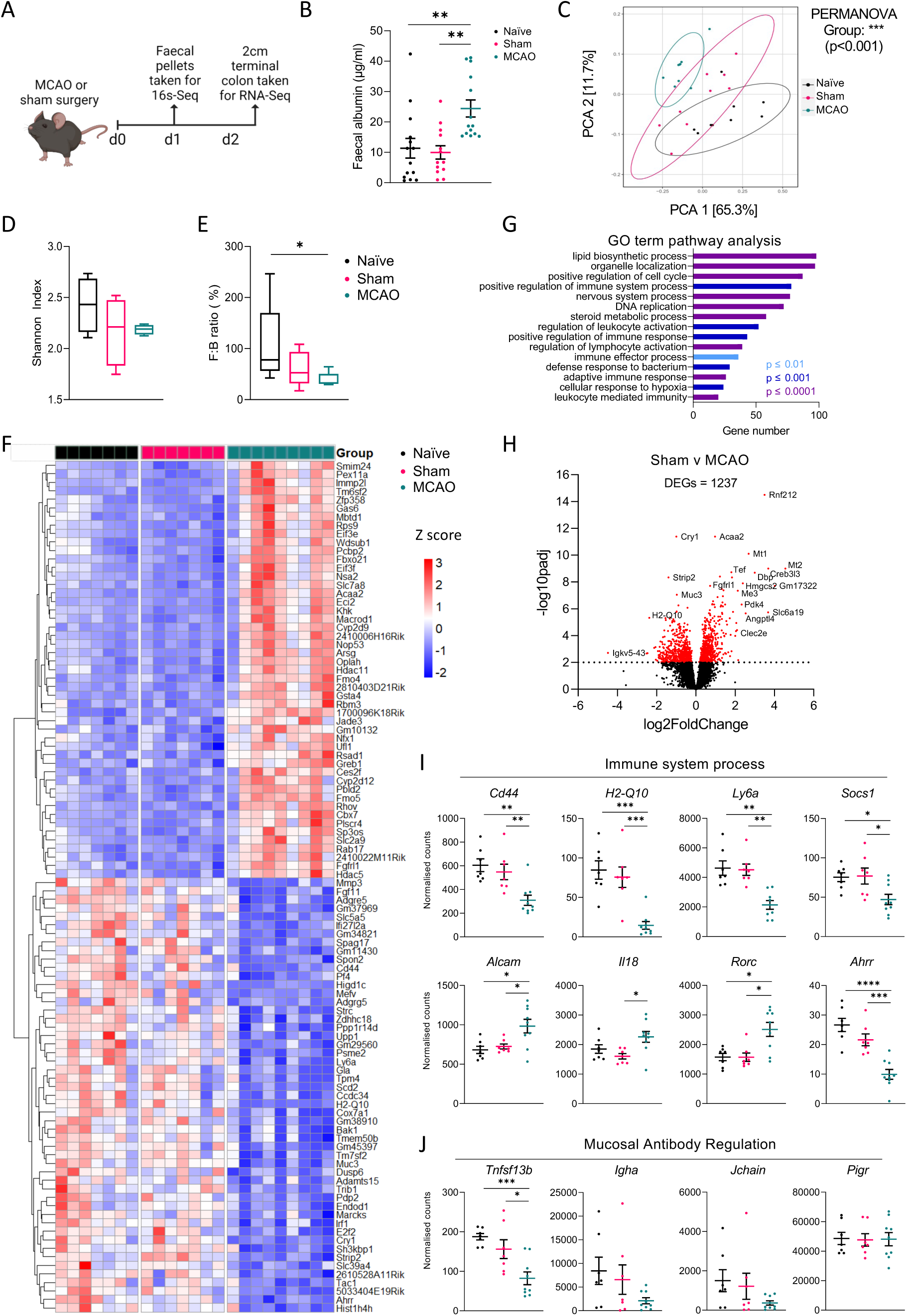
Stroke alters the intestinal transcriptome and microbiota composition. A) Schematic showing experiment design: C57BL/6 mice underwent MCAO, sham surgery, or no surgery (naïve). Faecal pellets were collected at 24h and mice euthanised at day 2, where colon tissue was harvested for bulk RNAseq. B) Faecal albumin concentration at 24h, data pooled from 3 independent experiments, n=13-14. (C-E) 16S rRNA sequencing of fecal microbiota from naïve, sham and MCAO mice. C) Bray-Curtis dissimilarity index, D) Shannon index & E) Ratio of *Firmicutes*:*Bacteroides* (F:B), data pooled from 2 independent experiments, n=8-9. (F-J) Bulk RNA sequencing of colonic tissue from naïve, sham and MCAO mice. F) Heat map of genes with 50 highest and 50 lowest differentially expressed genes (DEGs; p<0.01), expressed as z-scores. G) 15 curated pathways from GO-term pathway analysis. H) Volcano plot of top 1000 significant DEGs (p <0.01) from comparison of sham and MCAO groups. Positive fold change indicates increase following stroke, negative fold change indicates decrease following stroke. (I-J) Selected exemplar genes identified from GO term pathways including I) immune system process and J) mucosal antibody regulation. Data pooled from 2 independent experiments, n=7-9. Data presented as mean +/-SEM. Statistical tests: B, D, I+J – one-way ANOVA w/ Tukey post-hoc. C= PERMANOVA, * p <0.05, ** p <0.01, *** p <0.001, **** p <0.0001.

Next, to investigate changes in intestinal physiology following stroke we first performed bulk RNA-seq on colonic gut tissue (Figure 1F). This revealed ∼1200 highly differentially expressed genes (DEGs; p<0.01) between sham and stroke groups (Figure 1F-H), with comparable changes in gene expression between the intestines of naïve and stroke mice (Fig. S1C). In contrast very few genes were significantly different between naïve and sham mice (Fig. S1C). GO term enrichment highlighted pathways that were significantly altered by stroke when compared to sham controls, including “positive regulation of the immune system”, “leukocyte activation” and “defence response to bacterium” (Figure 1G), suggesting changes in the intestinal immune response may be associated with stroke. We further explored individual genes involved in these pathways (Figure 1I), and additionally identified a number of genes involved in humoral immunity following stroke (Figure 1J). Indeed, we noted that one of the most differentially expressed genes was *Igkv5-43*, a gene encoding an Immunoglobulin light chain, which decreased following stroke (Figure 1H). We also subsequently identified alterations in genes associated with B cell regulation and mucosal antibody production following stroke – including a significant reduction in *Tnfsf13b* (encoding BAFF) which supports B cell/plasma cell survival, and trends indicating potential changes in *Igha* and *Jchain* – genes associated with intestinal IgA secretion (Figure 1J). In addition, we noted significant disruption of circadian clock gene expression following stroke - including *Arntl* and *Nr1d1* (Fig. S1D) - transcriptional regulators that we have previously shown to be associated with regulation of the intestinal IgA response (Penny et al., 2022). Together these observations provoke the question as to whether stroke dysregulates homeostatic mucosal antibody responses to the microbiota.

### Intestinal B cell and IgA responses are altered following stroke

Our transcriptomic analyses of whole intestinal tissue were suggestive of altered B cell and IgA responses following stroke. Previous findings suggest stroke leads to acute suppression of B cell populations in peripheral organs such as the spleen (McCulloch et al., 2017), although whether this effect extends to mucosal barrier tissues and associated lymphoid tissues remains unclear. Thus, we focused our analyses on the major sites of IgA generation and secretion – namely the Peyer’s Patches (PP) and small intestinal lamina propria (Lycke and Bemark, 2012). Within 48 hours after surgery we observed a significant reduction in the frequency and number of germinal centre (GC) B cells (Figure 2A-B) and IgA^+^ class-switched GC B cells (Figure 2C-D) in the Peyer’s patches of MCAO mice, suggesting stroke may m GC activity in mucosal associated lymphoid structures. In contrast, we observed only a trend towards reduced frequencies of total and naïve IgD^+^ B cells in the Peyer’s Patches (Fig. S2A-B).

**Figure 2.**
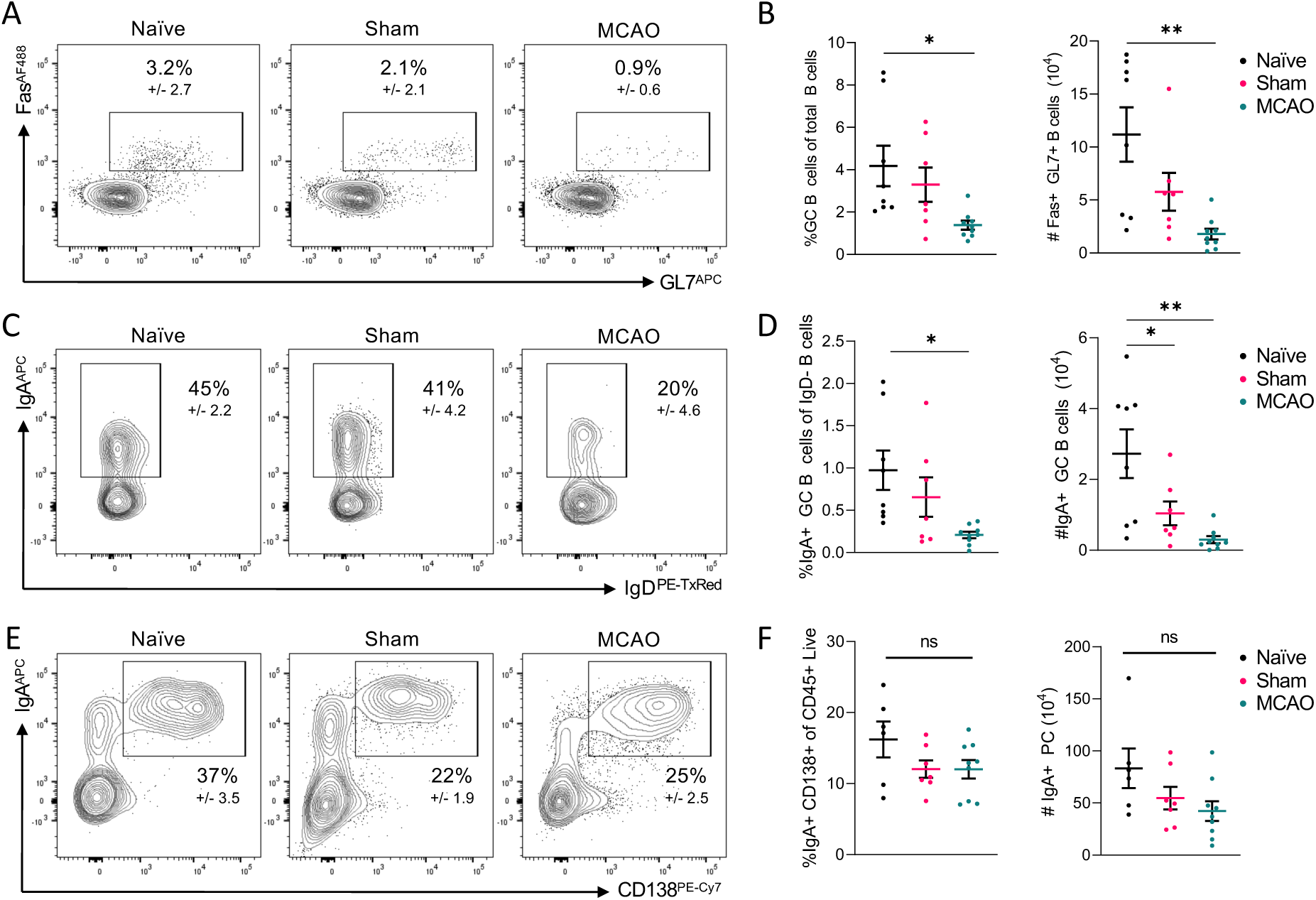
Stroke leads to dysregulation of the mucosal associated B cell compartment. (A-D) Flow cytometry analysis of Peyer’s patches B cell compartment. See also Supplemental Figure 2 for gating strategies. A) Representative flow plots of GL7+Fas+ Germinal Centre (GC) B cells including percent of parental gate (Live, CD45+ B220+ CD19+), B) quantification of GC B cells as a percentage of total B cells, and cell number. C) Representative flow plots of IgA+ GC B cells and D) quantification as a percentage of IgD-class switched B cells, and cell number. (E-F) Flow cytometry analysis of small intestinal lamina propria plasma cells. E) Representative flow plots and F) percentage and number of IgA+ plasma cells from the small intestine. All data were acquired at 48hrs following surgery. n=6-9, data pooled from 2 independent experiments. Data presented as mean +/-SEM. Statistical tests; B, D & F – one-way ANOVA w/ Tukey post-hoc, * p <0.05, ** p <0.01.

While IgA class switching takes place within the PP, the major source of IgA secreted into the intestinal lumen are lamina propria-resident IgA^+^ plasma cells. In contrast to newly generated IgA+ B cells in the Peyer’s patches, the CD138+ IgA+ plasma cell populations within the small intestinal and colon lamina propria was largely maintained post stroke with only a trend towards reduced cell numbers in MCAO mice at this time point (Figure 2 E-F, Fig. S2D). In addition, we did not observe major changes in small intestinal lamina propria T cell populations following stroke, including Th17, Treg or γ∂ T cells (Fig. S2E-H). Together, these findings therefore suggest differential effects of post-stroke immune suppression on the generation of B cell responses versus long-lived antibody secreting cells in the intestinal tract.

### B cells and IgA+ plasma cells are increased in the brain and meninges after stroke

Recent work has begun to demonstrate important functions for B cells and IgA+ plasma cells present within the CNS and meninges at both steady-state and during disease, with increasing evidence that meningeal IgA+ plasma Cells may originate in the gut (Brioschi et al., 2021; Clatworthy, 2021; Fitzpatrick et al., 2020; Fitzpatrick et al., 2024; Rojas et al., 2019). Thus, we asked whether the changes seen in B cells and IgA responses within intestinal associated tissue post-stroke might also lead to altered humoral immune cell numbers in the brain and meninges (Figure 3A). While B cells and IgA+ plasma cells were very rare within the brain parenchyma in both sham and stroke, there was an increase in the frequency and numbers of these cell types specifically in the stroke-impacted hemisphere when compared to the contralateral hemisphere (no infarct) (Fig. S3). Moreover, we found naïve IgD+ B cells to be present in the meninges of naïve animals in line with recent evidence (Brioschi et al., 2021; Fitzpatrick et al., 2024), which were increased in frequency and number in MCAO but not sham animals (Figure 3B-D). Similarly, IgA+ plasma cells were relatively rare in the meninges of naïve and sham mice but observed to increase following stroke (Figure 3E-G). Immunofluorescence imaging revealed both cell types to be largely enriched in the dural lymphatic sinuses following MCAO (Figure 3H,I). Together these findings suggest stroke not only perturbs humoral immunity within the gut but also within the brain’s borders.

**Figure 3.**
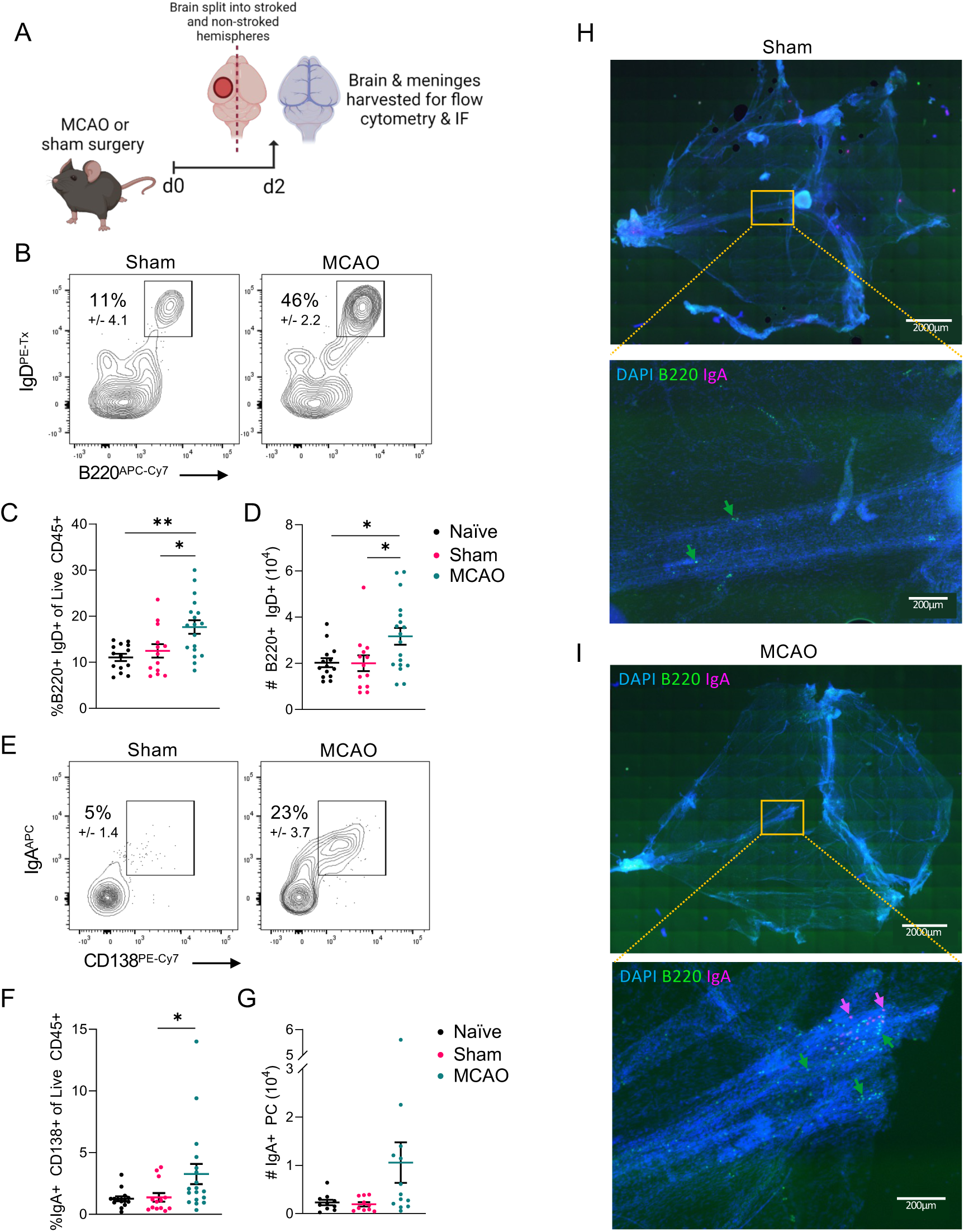
B cell and IgA+ plasma Cell numbers increase in the meninges of mice undergoing MCAO. A) Schematic showing experimental design: mice underwent MCAO and were allowed to recover for 48hrs before brains and meninges were harvested for downstream processing. See also Supplemental Figure 3. B) Representative flow plots, C) frequencies and D) numbers of naïve B cells (CD45+CD3- CD5- CD19+ B220^+^ IgD^+^) in the meninges. E) Representative flow plots, F) frequencies and G) numbers of IgA+ plasma cells (CD45+ CD3- CD5- CD19- B220- MHCII+ CD138^+^ IgA^+^) in the meninges. (H-I) Immunofluorescence staining for B220, IgA and DAPI and imaging of meninges from representative H) sham and I) MCAO mice. n=7-8, data pooled from 2 independent experiments. Data presented as mean +/-SEM. Statistical tests; B, D & F – one-way ANOVA w/ Tukey post-hoc, * p <0.05, ** p <0.01.

### Intestinal IgA dynamics differ in acute and chronic stroke

While IgA+ plasma cells numbers in the intestinal tract were largely stable at acute time points post stroke (Figure 2E-F), their capacity to secrete antibody can be dynamically regulated by a wide range of cues. We next aimed to clarify the effect of stroke on secretory IgA (sIgA) present within the intestinal tract itself. Surprisingly, we found a striking increase in faecal IgA levels following stroke in MCAO mice, but not in sham mice, as early as 24 hours following stroke (Figure 4A; normalised to total faecal protein Fig. S4A). IgA actively transported over the intestinal epithelium acts to bind directly to commensal microbes to modulate their colonisation, growth or biology. Thus, next we determined whether the stroke-induced increase in faecal IgA was associated with changes in the proportion of commensal bacteria bound by IgA (Figure 4B). Quantification of the global frequencies of IgA+ commensal microbes by bacterial flow cytometry revealed comparable proportions of bound bacteria 24hrs post-stroke (Figure 4C). We also tested for IgG binding, which can occur in the context of reduced barrier integrity - as observed following stroke (Figure 1B) - but observed little to no IgG labelling of bacteria following stroke (Fig. S4B). In contrast, a granular analysis of the relative proportions of commensal bacteria directly bound by IgA, or unbound by antibody, at the genera level by IgA-Seq revealed significant changes in IgA binding to a number of bacteria of the *Bacteroides* and *Firmicutes* genera (Figure 4D-E, Fig. S4C-D), suggesting IgA responses to these commensal bacteria could be altered post-stroke. Together these findings suggest that altered IgA responses and mucosal antibody labelling of commensal microbes are altered post-stroke.

**Figure 4.**
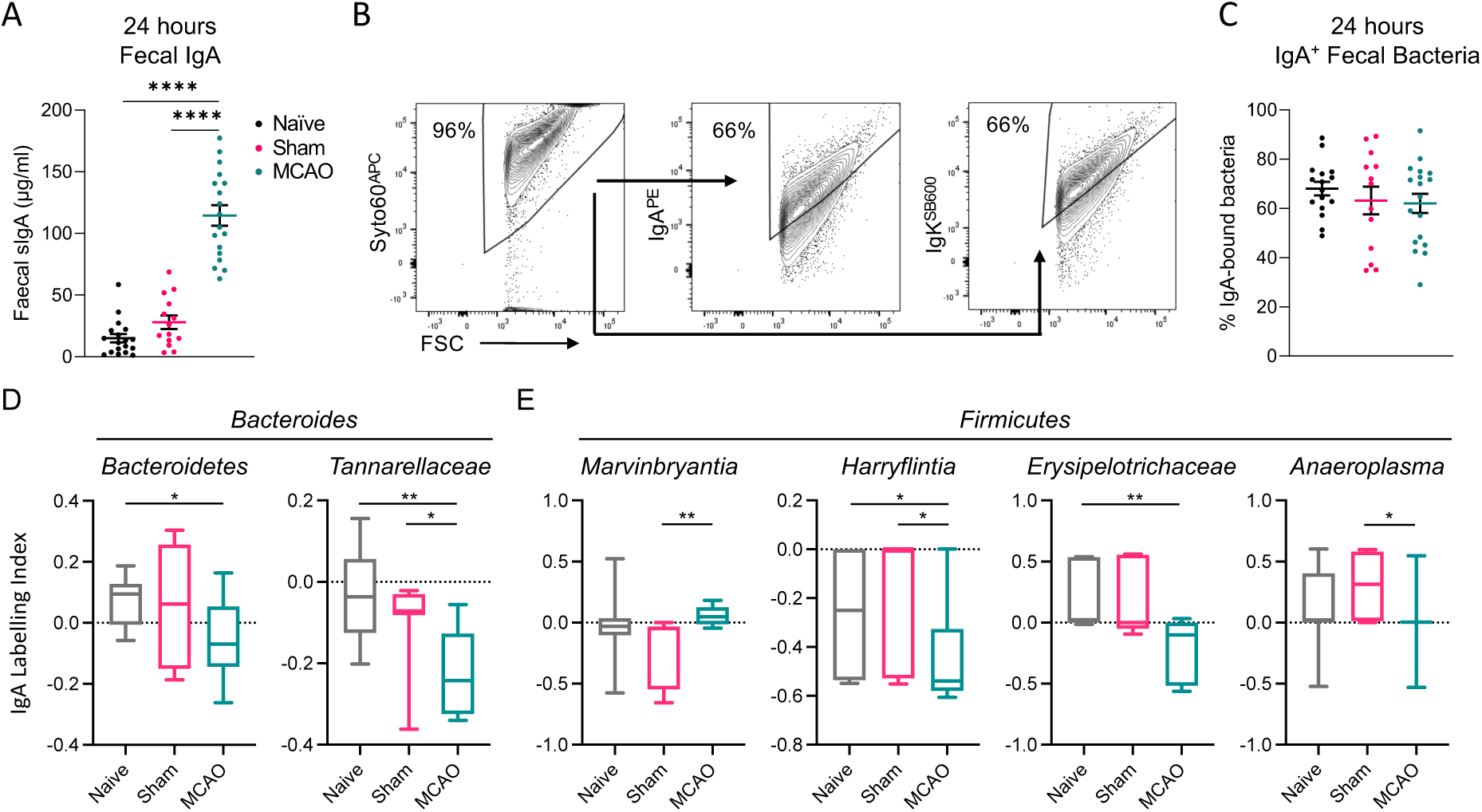
Intestinal IgA responses against the microbiota are perturbed following stroke. A) Faecal sIgA concentration at 24h, data pooled across 4 independent experiments, n=14-19. B) Gating strategy for bacterial flow cytometry. C) Percentage of IgA+ bacteria at 24h, data pooled across 4 independent experiments, n=13-18. (D-E) IgA-binding index (Kau Index) of D) *Bacteroides* and E) *Firmicutes* genera with a minimum abundance. Data pooled from 2 independent experiment, n=7-9 per group. Data presented as mean +/-SEM. Statistical tests; A – one-way ANOVA w/ Tukey post-hoc, D & E – Kruskal-Walis test with multiple comparisons, * p <0.05, ** p <0.01, *** p <0.001, **** p <0.0001.

### Intestinal sIgA responses are partially responsible for altered commensal microbiota

As changes to the microbiota were observed to occur rapidly following stroke (Figures 1+3 & Fig. S1), and we observed a spike in faecal IgA at acute time points following stroke (Figure 4A), we next asked whether increased luminal IgA could in part be responsible for stroke-induced alterations in the commensal microbiota. To investigate this we used IgMi mice, as reported previously (Penny et al., 2022; Waisman et al., 2007), which are unable to class-switch or secrete antibody but retain B cells and associated antibody-independent effector functions (Sahputra et al., 2018). Given IgA is the predominant antibody subtype in the intestines both in health - and immediately following stroke – this mouse model allowed us to investigate microbiome composition after stroke in the absence of intestinal IgA binding to luminal microbes (Figure 5A). IgMi mice and littermate controls (wild type; WT) were subject to sham surgery or MCAO. Importantly, the absence of secretory antibody had no impact on the initial infarct volume or stroke-associated acute behavioural deficits (Fig. S5A-C) – indicating acute stroke severity did not differ between IgMi and control mice subject to MCAO.

**Figure 5.**
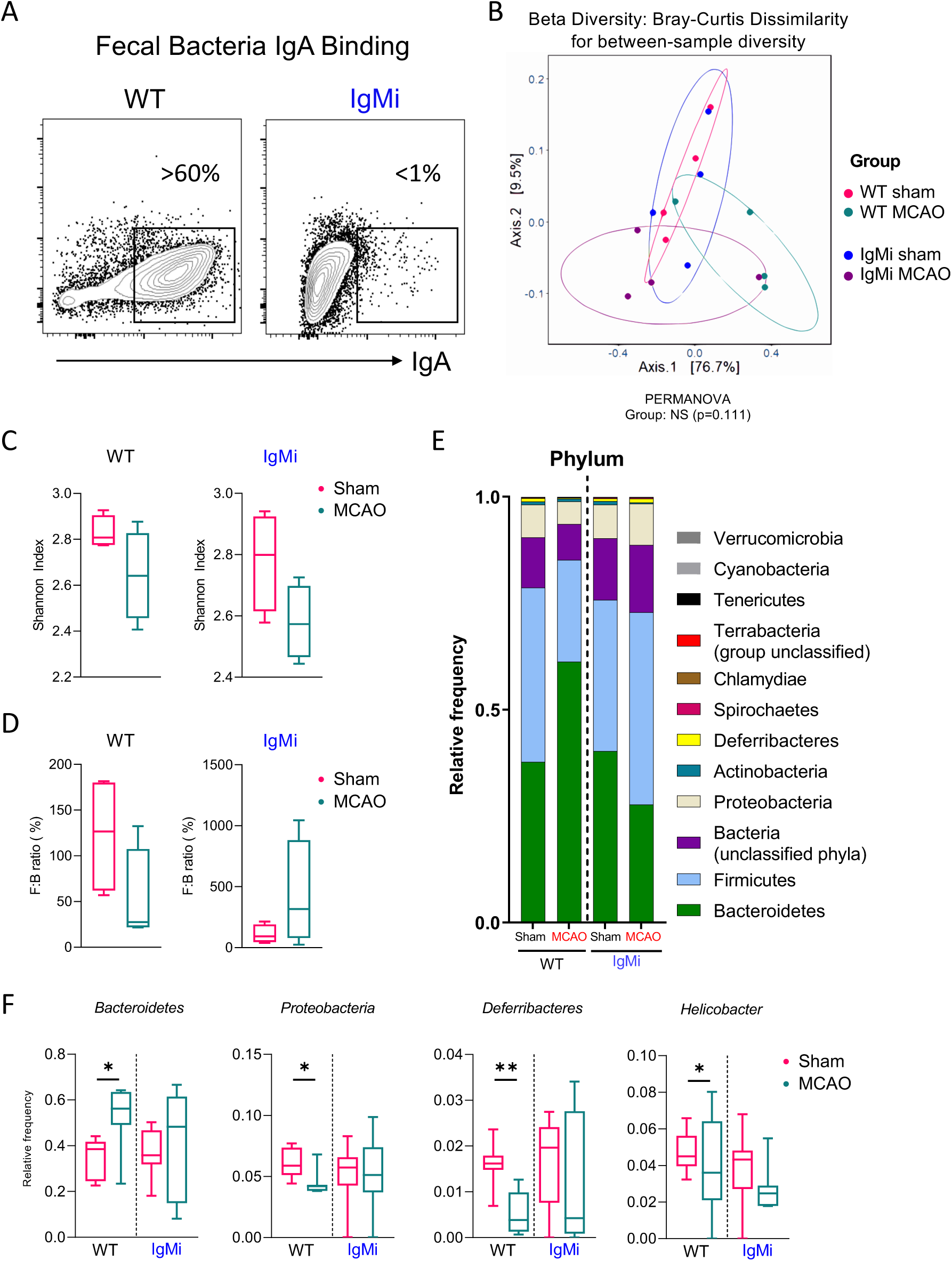
Stroke-associated alterations in microbiota composition are in part dependent on IgA. A) Representative flow cytometry plots showing IgA binding to bacteria isolated from faecal pellets of IgMi mice compared to littermate controls (WT). (B-F) shotgun metagenomic sequencing of fecal microbiota isolated from IgMi and WT mice 48h after undergoing sham or MCAO surgery. B) Bray-Curtis dissimilarity index, C) Shannon index & D) *Firmicutes*:*Bacteroides* (F:B) ratio, n=4. Data representative of 2 independent experiments. E) Stacked bar charts representing average composition of microbiota at Phylum level in WT and IgMi mice undergoing sham or MCAO. F) Select individual genera from sham and stroke WT and IgMi mice, data pooled from 2 independent experiments, n=7-8. Data presented as mean +/- SEM. Statistical tests; B = PERMANOVA, F = Kruskal-Wallis test with multiple coparisons, * p <0.05, ** p <0.01.

IgMi mice and littermate controls undergoing sham surgery had largely comparable global microbial diversity (Figure 5B), in line with our previous findings that lack of IgA does not disrupt global community composition at homeostasis (Penny et al., 2022). As expected, microbial communities segregated differentially from between stroke and sham groups in WT animals (Figure 5B). Notably however, while the microbial signature of IgMi mice also exhibited a clear change following stroke, IgMi MCAO mice exhibited differential clustering to WT MCAO mice - suggestive of potential differences in microbial composition and diversity following stroke in the presence or absence of secretory antibody (Figure 5B).

In line with our previous observations (Figure 1D), stroke led to a trend towards a reduction in Shannon diversity index when compared to sham controls, which occurred both in the presence and absence of IgA (Figure 5C). Again, we observed a reduction in the *Firmicutes* to *Bacteroides* ratio in MCAO WT mice (Figure 5D, Fig. S4F), as expected and in line with our prior observations (Figure 1E). In contrast, the reduced ratio between these two major commensal phyla was no longer evident following stroke in IgMi mice (Figure 5D+E). As we observed altered IgA-binding to *Firmicutes* and *Bacteroides* following stroke in wild type animals (Figure 4), this suggests perturbations in IgA responses following stroke may in part contribute to stroke-driven changes in intestinal microbial community diversity and composition.

Next, we further investigated the extent of stroke-induced changes to commensal microbiota composition in the presence or absence of IgA (Figure 5E-H, Fig. S5D-F). Our analysis revealed multiple specific genera that were altered in abundance by stroke in WT MCAO mice when compared to sham controls, but which were not significantly altered following stroke in IgMi mice which lack secretory IgA (Figure 5F). Despite these specific changes we also observed changes in microbial abundance that were observed in both genotypes (Fig. S5E), as well as stroke-induced changes that only reached significance in IgMi mice, and not WT mice, following stroke (Fig. S5F). Together these findings suggest changes in mucosal IgA responses may contribute in part to microbiota alterations observed in experimental models of stroke.

## Discussion

Alterations in gastrointestinal function and commensal microbiota composition following stroke are well described both in patients (Cui et al., 2023; Xia et al., 2019; Xu et al., 2019; Yamashiro et al., 2017; Yin et al., 2015) and animal models (Benakis et al., 2016; Brichacek et al., 2020; Houlden et al., 2016; Singh et al., 2016; Stanley et al., 2016; Stanley et al., 2018). Importantly, changes in microbial communities correlate with worsened stroke outcome and secondary complications (Benakis et al., 2016; Brichacek et al., 2020; Singh et al., 2016; Stanley et al., 2016; Tuz et al., 2022; Xia et al., 2019). Indeed, antibiotic intervention post-stroke can lead to improved outcomes (Benakis et al., 2020b; Liu et al., 2022). Despite these advances the mechanistic basis that underpins stroke-induced intestinal phenotypes and changes to commensal microbial communities remain poorly understood.

Recent findings indicate stroke induces dysregulation of multiple intestinal immune populations (Benakis et al., 2016; Brea et al., 2021; Diaz-Marugan et al., 2023; Ge et al., 2022; McCulloch et al., 2017; Oyama et al., 2018) . Moreover, intestinal tissue-resident immune cell networks are critical in regulating the commensal microbiota (Belkaid and Hand, 2014) – however whether maladapted immune responses to the microbiota mechanistically drive changes in the commensal community post stroke remains unclear. One major mechanism of host-commensal interactions is the production and transluminal secretion of IgA, which physically binds to bacteria to modulate their growth, survival, and broader biology (Bunker and Bendelac, 2018; Pabst and Slack, 2020; Rollenske et al., 2021). Here we demonstrate acute changes in IgA production following MCAO-induced stroke in mice. While the total frequency of the faecal bacteria bound by IgA did not change immediately following stroke, IgA-seq revealed discrete changes in IgA binding of a subset of *Bacteroides* and *Firmicutes.* In contrast, the ratio of these two major phyla was not affected after stroke in IgMi mice, which lack secretory IgA. Together these findings suggest that acute changes in IgA production and binding may in part contribute to post-stroke alterations in commensal communities. Despite these observations, other microbial changes occurred following stroke independent of the presence of mucosal antibody suggesting other mechanisms contribute to changes in the microbiota. Importantly IgMi mice exhibited lesions volumes and behavioural deficits equivalent to control littermates, suggesting IgA-dependent alterations of the microbiota do not determine the severity of stroke outcomes. Despite this we observed differences in the frequency of the microbiota bound by IgA at chronic time points, suggesting stroke may nonetheless have long-term consequences for host-commensal interactions, while the impact of these changes on the long-term prognosis of stroke remain unclear.

The mechanisms through which IgA responses are perturbed post-stroke remain to be elucidated. A prior seminal study demonstrated the loss of splenic marginal zone B cells in MCAO mice, driven by noradrenaline release and suppression of peripheral B cells via the β2-adrenergic receptor (McCulloch et al., 2017). In line with our findings, and that of a previous study (Oyama et al., 2018), we also detected reductions in germinal centre B cells within the intestinal-associated Peyer’s patches in MCAO mice, including reductions in IgA-class switched B cells (Figure 2) – suggesting similar mechanisms could suppress active B cell responses and class-switching within mucosal tissues as well. In contrast, IgA+ plasma cells – which reside within the intestinal lamina propria – were not significantly reduced in MCAO mice suggesting differential impacts of stroke on terminally differentiated antibody producing cells, or within tissue sites. While the reason for these differences requires further investigation, plasma cells are notably long-lived and non-proliferative cells – in contrast to lymphoid tissue B cells within active germinal centres – which may make them less susceptible to cell death induced via neuropeptide release. Alternatively, the quality or magnitude of innervation within plasma cell niches could differ, or those signals may elicit differential outcomes due to intrinsic differences in plasma cell biology. Moreover, while enhanced IgA levels in the faeces may reflect an increase in the magnitude of antibody production by tissue-resident plasma cells on a per cell basis, it is also feasible that increased gut leakiness noted post stroke negates the potentially rate-limiting step of active epithelial IgA transport by the polymeric immunoglobulin receptor (pIgR) - required when the epithelial barrier is intact - instead allowing for passive IgA diffusion into the lumen at greater concentrations. Future studies are needed to further define the neuroimmune crosstalk that ensues following stroke to define the extent of immunological change and understand their consequences.

Recent advances have demonstrated the presence of antibody secreting cells within the CNS and meninges in health and disease (Brioschi et al., 2021; Fitzpatrick et al., 2020; Probstel et al., 2020; Rojas et al., 2019). Intriguingly, these studies suggested IgA-producing cells found within the meninges were of intestinal origin (Fitzpatrick et al., 2020; Probstel et al., 2020; Rojas et al., 2019), indicating potential for gut-CNS migration, although more recent studies suggest dural-associated lymphoid tissues may also support local B cells responses (Fitzpatrick et al., 2024). Here we also observed B cells and IgA+ plasma cells present in the brain parenchyma and meninges in the acute stages following stroke – in line with previous findings that observed B cells within the brains of animals at chronic time points following stroke (Doyle et al., 2015; Ortega et al., 2020). Intriguingly, this coincided with a reduction of IgA+ B cells in the Peyer’s patches of the intestine raising the possibility of migration between these two sites post-stroke, although further studies with more elegant models will be required to disentangle the origins of IgA-producing cells following stroke.

Notably the absence of IgA in IgMi mice did not alter short term stroke severity, raising the question as to why humoral immune responses may expand or be recruited to the CNS in this context. Fitzpatrick et al demonstrated that meningeal IgA production could act as a firewall for circulating microbes with potential to enter the CNS. Stroke leads to a breakdown of intestinal barrier function in preclinical models and increases susceptibility to secondary infection. Indeed, bacteraemia is commonly reported following stroke in mice, with culturable microbes present in systemic organs including the liver, lungs and spleen (Diaz-Marugan et al., 2023; McCulloch et al., 2017; Stanley et al., 2016). Thus, it is attractive to hypothesise that an increase in antibody producing cells within the meninges post stroke may help protect the brain from translocating or invading bacteria that have entered the bloodstream.

Together this study highlights the need for further investigation into the role of the gut-brain axis and role of the intestinal immune system in the context of the stroke, and has the potential to open new therapeutic avenues for the management of both the primary pathology and secondary complications arising in patients.

## Methods

### Animals

All C57BL/6 mice were sourced from Charles River. IgMi mice were provided by Prof Ari Waisman (University of Mainz, Germany). Transgenic lines were bred in-house at the Biological Services Facility (BSF) at the University of Manchester. All mice were maintained under specific pathogen-free conditions and kept in sterilised filter-topped cages under a 12-hour light/dark cycle. Mice had access to food and water *ad libitum*, unless otherwise indicated. Mice were acclimatised for a period of at least 7 days before any procedures were performed. All animal experiments were conducted in accordance with the regulations stipulated in the Animal (Scientific Procedures) Act 1986. The procedures conducted for this research project were performed by individuals holding UK personal licences under the project licence issued to Prof Stuart Allan (P28AA2253, Jul 2017-2022). Protocols for experimental work were reviewed and approved by the University of Manchester Animal Welfare and Ethical Review Board.

### Experimental design

The ARRIVE guidelines were consulted during experimental design (Percie du Sert et al., 2020). All experiments were performed in mice aged between 7-19 weeks of age. Stroke experiments were conducted using male mice only. For experiments using transgenic lines, age-matched littermate controls were used, unless otherwise specified. Mice were randomised to intervention groups using block randomisation, to avoid cage effects. Exclusion criteria included animals that died under procedure or where welfare declined such that mice would need to be immediately culled. Mice that were suspected not to have had a stroke were also excluded based on front limb symmetry by tail suspension, and/or absence of brain lesion as shown by MRI or Cresyl violet staining. Following stroke or sham surgery, mice were housed with others of the same group (i.e. stroke mice were only housed with other stroke mice) to avoid post-surgery transfer of microbiota. Blinding to experimental groups and/or genotypes was only done where specified (i.e. neuroscore analysis) due to need to track and separate genotypes during surgery. Mice were euthanized in accordance with Home Office regulations, either by exposure to a rising concentration of CO_2_, cervical dislocation or transcardial perfusion of saline under anaesthesia.

### Stroke surgery

A modified version of the transient MCAO protocol was used to induce experimental cerebral ischaemia (Longa et al., 1989). A small incision was made in the scalp and a Doppler probe was attached using VetbondTM to allow monitoring of cerebral blood flow. A midline incision to the throat was made and soft tissues pulled apart to expose the left common carotid artery which was dissected free from surrounding nerves. Two ligatures were made around the left common carotid artery (spaced approximately 3mm apart) before it bifurcates into the left internal and external carotid arteries, which were then also isolated and tied off. A small incision was made in the common carotid artery in the region between the two ligatures, then a nylon monofilament (diameter 180µm) coated with silicon was inserted into the common carotid artery, advanced into the middle cerebral artery via the internal carotid artery by 10mm, and left in place for 30 minutes (this step was omitted for sham surgeries). Successful occlusion of the middle cerebral artery was confirmed by a consistent drop in cerebral blood flow as measured by Doppler. The filament was then withdrawn and the wound closed with 6-0 Vicryl sutures (Ethicon). Post-operative care was given as follows: Buprenorphine at 0.05mg/kg and 500µl saline was injected subcutaneously for analgesia and fluid replacement respectively, and the animals were placed in a heated cabinet during recovery. Mice were provided with floor-fed wet mash and given 0.05mg/kg Buprenorphine and 300µl saline daily until sacrifice.

Post-stroke neurological deficits were tested at day 1 after surgery using a 28-point scale adapted from Clark et al. (1998). Seven criteria were assessed; in brief, these were body asymmetry, gait, climbing ability, circling behaviour, front limb asymmetry, compulsory circling and response to whisker touch. Neurological assessment was carried out by an experienced researcher who was blinded to experimental groups. To accurately quantify ischaemic lesion volume, mice were scanned by T2-weighted MRI at day 1 post-surgery. MRI scans were acquired using a 7T horizontal bore magnet (Agilent Technologies, CA, USA) connected to a BrukerAvance III console (Bruker Biospin, MA, USA) using a surface transmit-receive coil. Mice underwent a short localisation scan, followed by a 19-minute T2 acquisition in which 20 coronal scans of 1mm depth were acquired. Lesion volumes were measured in Fiji (ImageJ, NIH) by calculating the oedema ratio (area of contralateral hemisphere/ipsilateral) and then multiplying this by the lesion area to give an oedema-corrected lesion volume (Nouraee et al., 2019).

### Whole mount meninges staining

Mice were transcardially perfused with sterile saline under anaesthesia. Skull caps with the intact dural meninges were collected and incubated in 4% PFA at 4°C for 6 hours. The meninges were then carefully removed from the interior surface of the skull and transferred to a 24-well plate for staining with the following: fluorescent-conjugated antibodies AF488-B220, PE-CD138 and APC-IgA in 2% BSA 0.2% Triton and 2% donkey serum in PBS. Meninges were incubated in the dark on a shaker overnight at 4°C, followed by washing with PBS then DAPI was added for 5 min. Meninges were washed again with PBS before mounting. To mount, meninges were floated in a petri dish containing PBS and carefully mounted onto SuperFrostTM Plus slides (Thermo Fisher Scientific) and left to dry for 1-2 hours. Once dry 1-2 drops of ProLongTM Gold antifade mountant (Invitrogen) were added and slides were coverslipped and allowed to dry in the dark before imaging. Whole-mount images were collected on an Olympus BX63 upright microscope using a 10x/0.4 UApo/340 objective and captured and white-balanced using a DP80 camera (Olympus) in colour mode through CellSens Dimension v1.16 (Olympus). Specific band pass filter sets for DAPI, FITC, Texas red and CY5 were used to prevent bleed through from one channel to the next. Images were then processed and analysed using Fiji (ImageJ, NIH).

### Faecal pellet collection

Fresh faecal pellets were obtained either by placing mice in clean cages without bedding and waiting for defecation or removed directly from the colon at necropsy. In each case care was taken to ensure collection tools were sterilised between mice and no contamination of samples occurred. To separate the bacteria and supernatants for ELISA, faecal pellets were resuspended in sterile PBS at a concentration of 1µl/mg and incubated on ice for 20min. Pellets were homogenised for 30 seconds at 4.0m/s on a tissue homogeniser (Fastprep 24, MP Biomedical) then centrifuged for 5 min at 200g to remove debris. Supernatants were filtered through a 70µm cell strainer and centrifuged again at 8000g for 5min to pellet the bacteria. Supernatant and bacterial pellets were stored at -80°C until analysis.

### ELISA

Faecal albumin and IgA were quantified in normalised faecal supernatants using either ELISA Quantitation Sets (Bethyl Laboratories) or Mouse IgA ELISA kits (Invitrogen) following manufacturer’s instructions.

### IgA ELISpot

An IgA ELISpot (Mabtech, Sweden) was conducted on single cell suspensions from the small intestine. PVDF type MSIP plates (Merck, Germany) were pre-treated with 15µl 35% ethanol for <1min and coated with IgA mAB MT45a overnight in the fridge. Plates were washed and blocked with 10% FCS for 30 minutes at room temperature. Blocking solution was removed, 100k cells were added to each well and the plate was covered and incubated at 37°C with 5% CO2 for 16-24 hours. The plate was then washed 5 times with PBS and detection MT39A-biotin was added and the plate was incubated for 2 hours at room temperature. The plate was washed again, streptavidin-ALP solution was added and the plate was incubated for 1 hour at room temperature. After washing the plate again, substrate solution was added until distinct spots emerged and the reaction was stopped by washing excessively with tap water. The plate was allowed to dry and spots were then counted on an AID ELISpot reader (Biosys, USA).

### Bicinchoninic acid (BCA) assay

To measure total protein in faecal pellet supernatants, the Pierce™ BCA Protein Assay Kit (ThermoFisher Scientific) was used according to manufacturer’s instructions. In brief, standards and sample replicates were pipetted onto a 96-well plate, after which a working solution was added at a sample:solution ratio of 1:20. Plates were then covered and incubated at 37°C for 30 min before cooling and reading absorbance on a spectrophotometer (Versamax, Molecular Devices) at an optical density of 562nm.

### Microbiome sequencing

Whole faecal pellets were obtained as described above. Pellets were collected either into sterile tubes on ice and frozen at -20°C until DNA extraction (for 16-seq by the Centre for Genomic Research (CGR), Liverpool, UK), or directly into collection tubes containing DNA stabilisation buffer for sequencing by Transnetyx (Cordova, TN, USA). For sequencing carried out by the CGR, bacterial genomic DNA was isolated with the DNeasy PowerLyzer PowerSoil kit (Qiagen) following manufacturer’s instructions. Briefly, faecal pellets were homogenized and bacteria isolated as described above. The bacterial pellet was then transferred to a PowerBead Tube and homogenized again at 4m/s for 30 sec. Protein was precipitated and inhibitors were removed before loading the sample into an MB Spin Column to bind the DNA before eluting into an Eppendorf. All steps were done in a tissue culture hood with sterile equipment and unopened kit reagents to avoid contamination of samples.

### Pre-amplification

Pre-amplification of the V3/V4 region of 16S rRNA was performed by PCR in triplicate using 2xKAPA HiFi Hot Start ReadyMix (Roche) using primer pairs containing adaptor sequences for down-stream use on Illumina platforms, as follows:

16S Amplicon PCR Forward Primer:

5’ TCGTCGGCAGCGTCAGATGTGTATAAGAGACAGCCTACGGGNGGCWGCAG

3’

16S Amplicon PCR Reverse Primer:

5’ GTCTCGTGGGCTCGGAGATGTGTATAAGAGACAGGACTACHVGGGTATCTAAT CC 3’

Prior to amplification of all samples, a subset of samples underwent amplification with either 10, 15 or 20 cycles/ rounds of amplification and an aliquot was run on a 1.5% agarose gel to ensure the PCR product was of the expected length of ∼380bp. Products of the correct length were formed at 10 cycles, thus all the samples were amplified at this number of cycles. First round PCR products were shipped on dry ice to the CGR for downstream sequencing and analysis.

### Bacterial flow cytometry & IgA-seq

Samples for bacterial flow cytometry and IgA-seq were processed as follows: bacteria from faecal pellets were isolated as described above. The bacterial pellet was then either processed for bacterial flow cytometry or processed for IgA-seq. For flow cytometry, pellets were stained with PE-IgA and SB600-IgK at 1:200 and SYTO-60 at 1:600 in FACS buffer for 30 min at 4°C. Samples were washed, resuspended in PBS and acquired on a BD LSRFortessaTM using DIVA software (BD Biosciences). Flow cytometry data was analysed and flow plots were produced in FlowJo V10.

For IgA-seq, pellets were stained as above After washing pellets were resuspended in PBS and a 200μl aliquot was taken and frozen at -80°C (total bacteria sample) and also a 50μl sample for flow cytometry analysis. The remaining bacteria were incubated with anti-PE beads (Miltenyi) for 30 min at 4°C before centrifugation and washing with FACS buffer. The sample was then passed through an LS column (Miltenyi) placed on a QuadroMACSTM Separator (Miltenyi) to enable binding of the IgA+ labelled bacteria to bind to the column (magnetic cell sorting, MACS). The IgA-fraction was collected and then the bound fraction was removed from the magnetic field and eluted to form the IgA+ fraction. 200μl samples were taken for flow cytometry analysis and the rest of the sample was pelleted frozen at -80°C until processed for genomic DNA extraction. Efficacy of MACS sorting was monitored by analysing the aliquots taken for flow cytometry.

The total bacterial samples and MACS sorted samples (IgA+ and IgA-fractions) underwent DNA extraction with the DNeasy PowerLyzer PowerSoil kit (Qiagen) as described above (2.12.4) and were then pre-amplified with primers targeting the V3/V4 region of 16S RNA with adaptor sequences for Illumina platforms as described in 2.8.3. Data were prefiltered in excel then analysed using R studio and GraphPad. IgA-binding index known as the Kau index (Kau et al., 2015) was calculated for each taxon present in an individual sample, with a mean and p-value calculated for each group. This gives a value between -1 and 1, indicating that a taxa is detected exclusively in the IgA-fraction or IgA+ fraction respectively, giving a relative measure of targeting that corresponds to a taxon’s overall abundance within the faecal sample, allowing for cross-group comparisons.

### Shotgun metagenomic sequencing

Shotgun sequencing was carried out by Transnetyx. In brief 2-3 faecal pellets, or caecal contents, were transferred to barcoded collection tubes containing DNA stabilisation buffer provided by Transnetyx. Samples were shipped to Transnetyx where DNA extraction was performed using the Qiagen DNeasy 96 PowerSoil QIAcube HT extraction kit. Library assembly was done using the KAPA hyperplus library preparation protocol, and sequenced with the Illumina NovaSeq instrument and protocol at a depth of 2 million (2x150 bp pair reads). Unique dual indexed (UDI) adapters were used to ensure that reads and/or organisms were not mis-assigned. Raw data (FASTQ files) were analyzed using the One Codex database, before exporting and analyzing in R Studio.

### Analysis of microbiome sequences

FASTQ sequences were uploaded to the One Codex database of whole microbial reference genomes and aligned using k-mers, where k=31. Based on the relative frequency of unique k-mers in the sample, probable sequencing or reference genome artifacts were filtered out of the sample. Then, relative abundance of each species was estimated based on the depth and coverage of sequencing across every available reference genome. Relative abundances were used to create stacked bar graphs of phyla and genera. PCoA were produced in R Studio. Only genera forming >1% of total abundance were used for generating bar graphs; genera with abundance below this are shown as “other”.

## Murine Transcriptional analysis

### RNA extraction

Tissue sections approximately 2cm long were dissected from the distal colon and flash frozen on dry ice. Samples were homogenized in 40µl of RLT buffer in lysing matrix tubes (MP Biomedical) for 30 sec at 4.0m/s on a tissue homogeniser (Fastprep 24, MP Biomedical). Total RNA was extracted using either an RNeasy mini kit (Qiagen) or the Norgen Single Cell RNA Purification Kit (Norgen Biotek Corp, Canada) following the manufacturer’s instructions. Sample RNA concentration was measured either using a Nanodrop for Real-Time PCR (qPCR), or using the QubitTM RNA HS Assay Kit (Thermo Fisher Scientific) and protocol for bulk-RNA sequencing.

### Bulk RNA-seq

Small intestine and colon RNA samples were frozen and shipped on dry ice to Novogene (Cambridge, UK) for sequencing. Novogene carried out in-house quality control and normalisation of raw FASTQ files. Data was analysed using the DESeq package in R Studio. Principal component analysis (PCA) plots were generated in R Studio. Z-scores were calculated and plotted as heatmaps in R studio. Fold change was calculated and used to generate volcano plots in GraphPad Prism v8. GO term analysis was carried out using the Gene Ontology Resource (Ashburner et al., 2000; Gene Ontology Consortium, 2021).

### Intestinal lamina propria isolation

To isolate the lamina propria lymphocytes, the small intestine and colon were dissected, associated fat removed and the tissue was cut open longitudinally, washed in PBS to remove luminal contents, and placed in 15ml PBS on ice. To fully remove luminal contents, tissue was vortexed and washed with PBS 3 times. Tissue was then placed in 15ml of stripping buffer (1mM EDTA, 1mM DTT and 5% FCS) for 10 min at 37°C on a tube rotator (Fisher Scientific), followed by a further 20 min incubation with rotation in fresh strip buffer to removed epithelial cells and intra-epithelial lymphocytes. The remaining tissue was washed with PBS and then incubated in 10ml of digestion buffer (0.1mg/ml collagenase/dispase (Roche) OR 1mg/ml collagenase D (Roche, Switzerland), and 20µg/ml DNase I (Sigma-Aldrich, St Louis, USA) for 45 min at 37°C on a tube rotator (Fisher Scientific. The resulting tissue and supernatant were passed through a 70µm cell strainer (Corning, New York, USA) and centrifuged to isolate lamina propria lymphocytes. Cell counts were obtained using a Casy TT counter (Roche Innovatis, Germany).

### Peyer’s patch processing

Between 4-6 Peyer’s patches were excised from the small intestine at necropsy and placed directly in digestion media containing 0.16mg/ml Liberase (Roche, Switzerland) and 40μg/ml DNase (Sigma-Aldrich, St Louis, USA) on ice. Peyer’s patches were then incubated in the digestion media at 37°C for 30 minutes before mashing through a 70μm cell strainer (Corning, New York, USA) into complete media.

### Brain and meninges cell isolation

Whole brains were placed in RPMI 1640 containing 2mM L-glutamine, 100 units/ml penicillin, 100µg/ml streptomycin, 10% foetal calf serum (FCS)), collagenase D (10µl/ml) and DNase (10µl/ml)) and cut finely with scissors before incubating for 30 min at 37°C with agitation. Resulting suspensions were filtered through a 70µm cell strainer (Corning, New York, USA) and centrifuged. The supernatant was discarded, the pellet resuspended, and centrifuged on a Percoll gradient. The uppermost layer containing myelin was carefully removed with a Pasteur pipette and remaining suspension diluted and mixed thoroughly in media, then centrifuged to isolate the leukocytes.

The meninges were peeled from the inside of the skull and placed in RPMI 1640 containing 500 U/ml DNase I and 140 U/ml Collagenase 8 (Sigma Aldrich) and incubated for 20 minutes at 37°C with agitation. The resulting suspension was filtered through a 70µl cell strainer (Corning, New York, USA) and centrifuged.

### Flow cytometry

Single cell suspensions were resuspended in FACS buffer (PBS with 4% FCS and 1mM EDTA) containing a surface stain antibody cocktail and incubated for 30 min at 4°C in the dark. To enable detection of transcription factors and cytokines, cells were resuspended in FoxP3 fix/perm buffer (eBioscience) for 30 min then centrifuged and washed with FoxP3 1x perm buffer (eBioscience). Intracellular antibody cocktails were made up in FoxP3 1x perm buffer and applied to cells for 30 min at 4°C in the dark. Fluorescence minus one (FMO) controls were used to help with gating for cytokines. All antibodies used for flow cytometry are listed in Table 2.1. After staining, samples were washed, resuspended in FACS buffer and acquired on a BD LSRFortessaTM using DIVA software (BD Biosciences). Flow cytometry data was analysed and flow plots were produced in FlowJo V10.

## Data Analysis and Statistics

All raw data were processed using Microsoft Excel and analysis was carried out in GraphPad Prism, unless otherwise specified. All flow cytometry data was analysed and flow plots were produced in FlowJo V10. Imaging data was processed and analysed using ImageJ (NIH). Experimental schematics were produced with Biorender. Data are reported as mean ± SEM unless otherwise stated, with details of specific statistical tests indicated in figure legends. For all data, p<0.05 = *; p<0.01 = **; p<0.001 = ***; p<0.0001 = ****.

## Acknowledgements

The authors acknowledge members of the Hepworth lab for critical discussion. Gareth Howell, David Chapman and the University of Manchester flow cytometry core for support. A.W. is supported by funding from the Deutsche Forschungsgemeinschaft (DFG) Project number 318346496 – SFB1292/2. Research in the Hepworth Laboratory is supported by a Sir Henry Dale Fellowship jointly funded by the Wellcome Trust and the Royal Society (Grant Number 105644/Z/14/Z), a BBSRC responsive mode grant (BB/T014482/1) and a Lister Institute of Preventative Medicine Prize.

## Author contributions

MH – Conceptualization, investigation, validation, formal analysis, data curation, writing (original draft), visualization. RW, LD-M, BDL – Investigation and validation. AW – Resources and methodology. SMA, CBL, DB, CB – Resources, methodology, writing (review & editing), supervision, project administration. MRH - Conceptualization, investigation, writing (original draft, review & editing), supervision, project administration and Funding acquisition.

---

**Figure S1.**
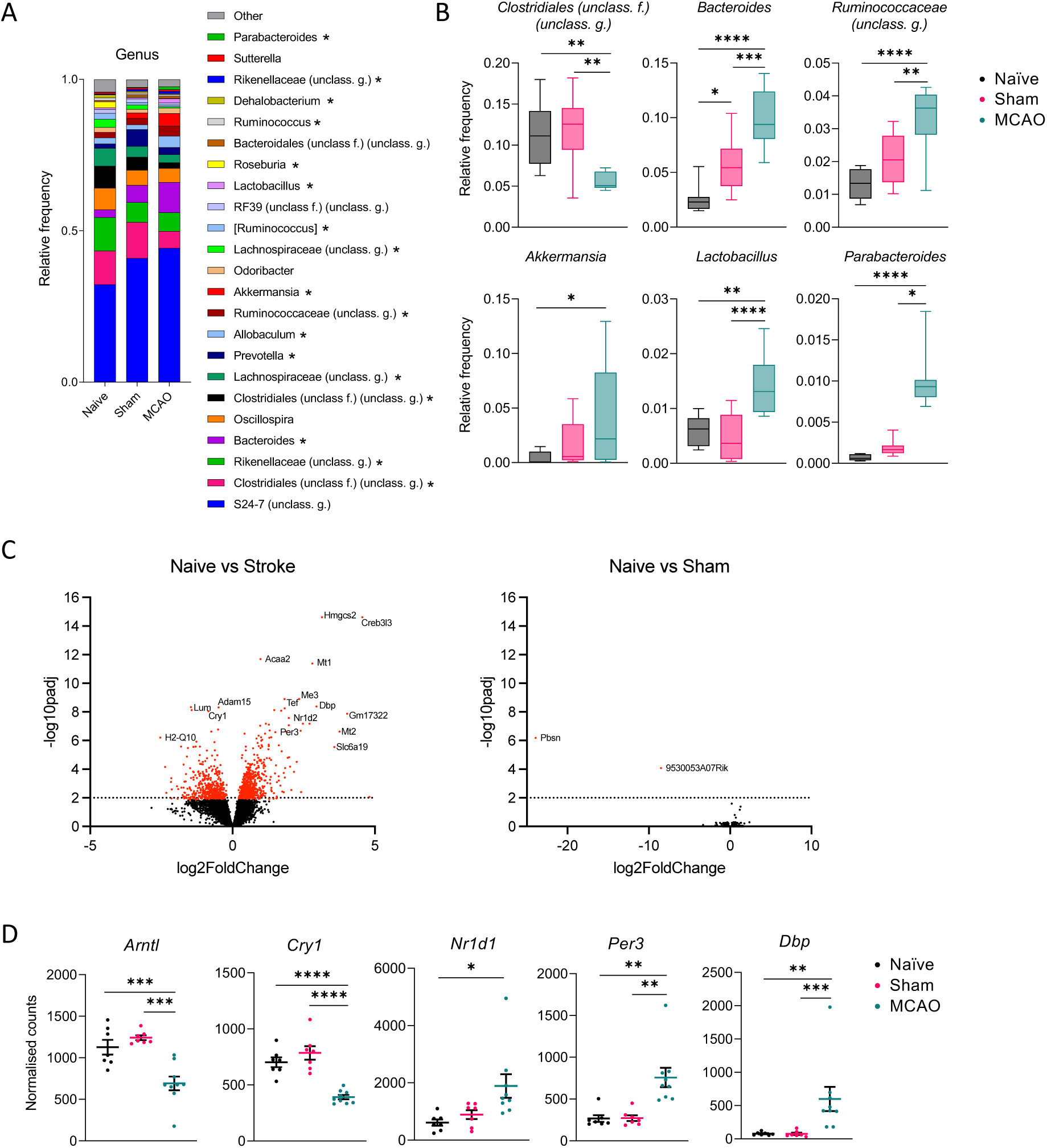
Effects of stroke on microbiota composition and intestinal gene signatures. Related to Figure 1. 16S rRNA sequencing of fecal microbiota from naïve, sham and MCAO mice. A) average faecal microbial composition at genera level. Data pooled from 2 independent experiments, n=8-9. B) Select bacterial genera identified to significantly alter in abundance following stroke. (C-D) Bulk RNA sequencing of colonic tissue from naïve, sham and MCAO mice. C) Volcano plots of significant DEGs from comparison of naive and MCAO groups (left panel) or naïve and sham groups (right panel). D) Selected exemplar genes identified from GO term pathways related to circadian rhythms. Data pooled from 2 independent experiments, n=7-9. Data presented as mean +/-SEM. Statistical tests; B & D one-way ANOVA w/ Tukey post-hoc, * p <0.05, ** p <0.01, *** p <0.001, **** p <0.0001.

**Figure S2.**
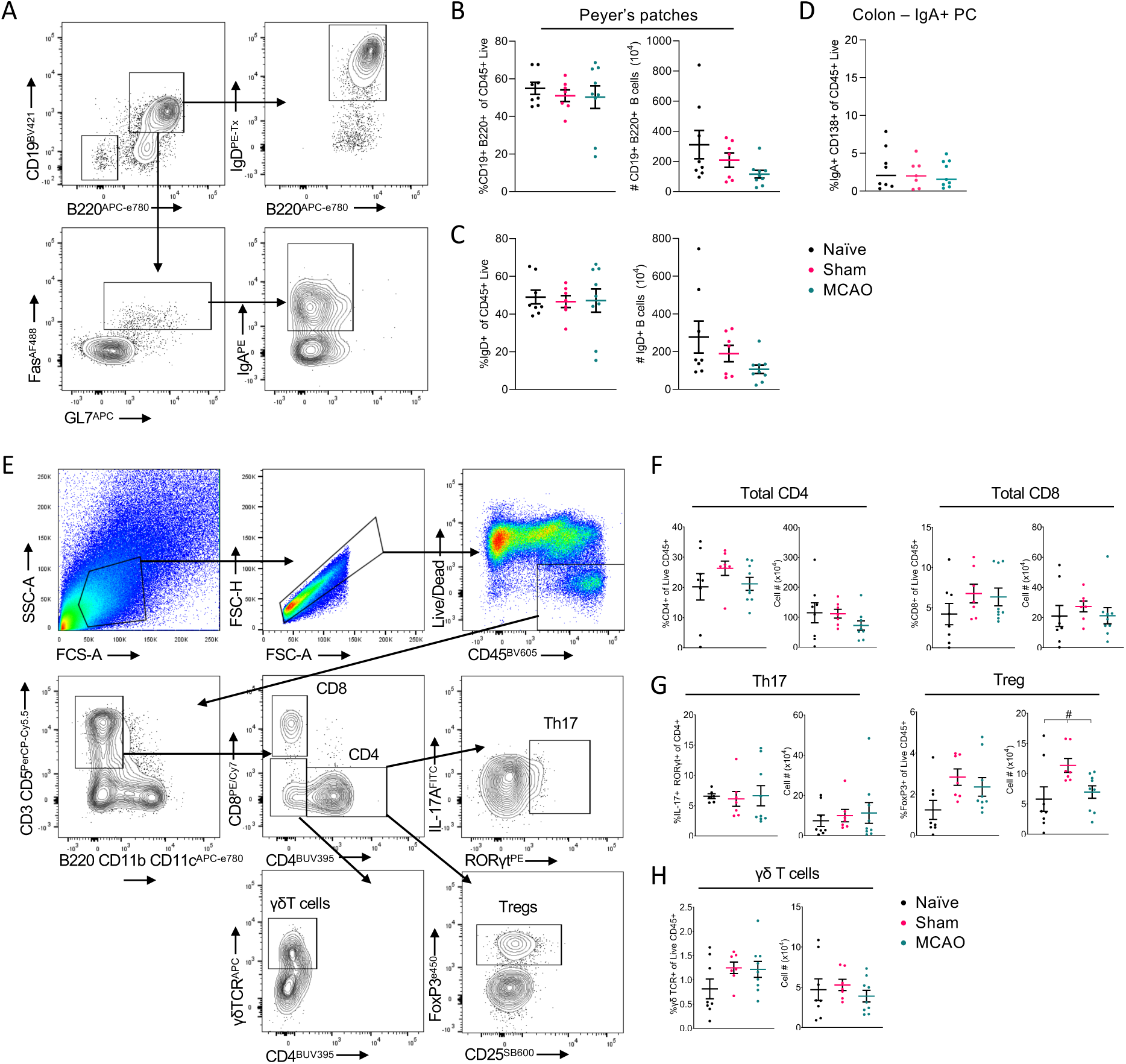
Immune profiling of the intestine and gut associated lymphoid tissue following stroke. Related to Figure 2. A) Exemplar gating strategy utilized for analysis of B cell compartment in Peyer’s patches. (B-C) Quantification of frequencies and numbers, of B) total B cells and C) IgD+ naïve B cells in Peyer’s patches, and D) quantification of frequencies of IgA+ plasma cells in the colonic lamina propria of naïve mice or mice undergoing sham or MCAO surgery. (E-H) Broader immunophenotyping of the small intestinal lamina propria following stroke. E) Exemplar gating strategies. (F-H) Quantification of frequencies and numbers of F) total CD4+ and CD8+ T cells, G) Tissue-resident CD4+ T helper cell subsets (RORγt+ Th17 and FoxP3+ Treg), H) γ∂ T cells. Data pooled from 2 independent experiments, n=7-9. Data presented as mean +/-SEM. Statistical tests; B, D, F-H – one-way ANOVA w/ Tukey post-hoc,

**Figure S3.**
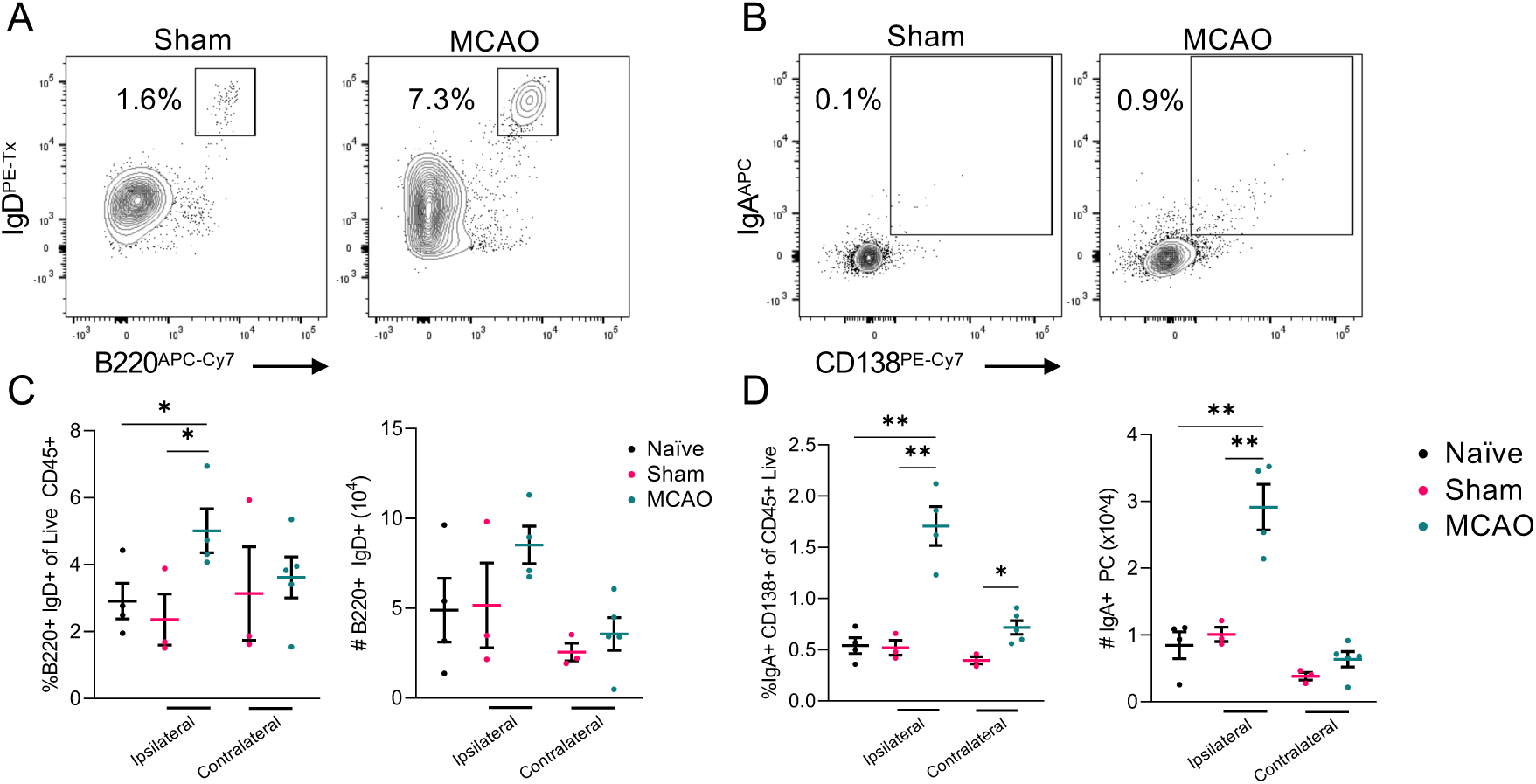
B cell and IgA+ plasma Cell numbers are increased in the ischaemic ipsilateral hemisphere of MCAO mice. Related to Figure 3. A) Representative flow plots of naïve B cells (B220^+^ IgD^+^) and B) IgA+ plasma cells (CD138^+^ IgA^+^) in the stroked hemisphere (ipsilateral) and non-lesional hemisphere (contralateral) of the brain. Gated as in Figure 2 and Figure S2 (C-D) Quantification of frequencies and numbers of C) naïve B cells (B220^+^ IgD^+^) and D) IgA+ plasma cells (CD138^+^ IgA^+^) in indicated brain hemispheres, n=3-5. Data representative of two independent experiments. Statistical tests; C-D – one-way ANOVA w/ Tukey post-hoc,

**Figure S4.**
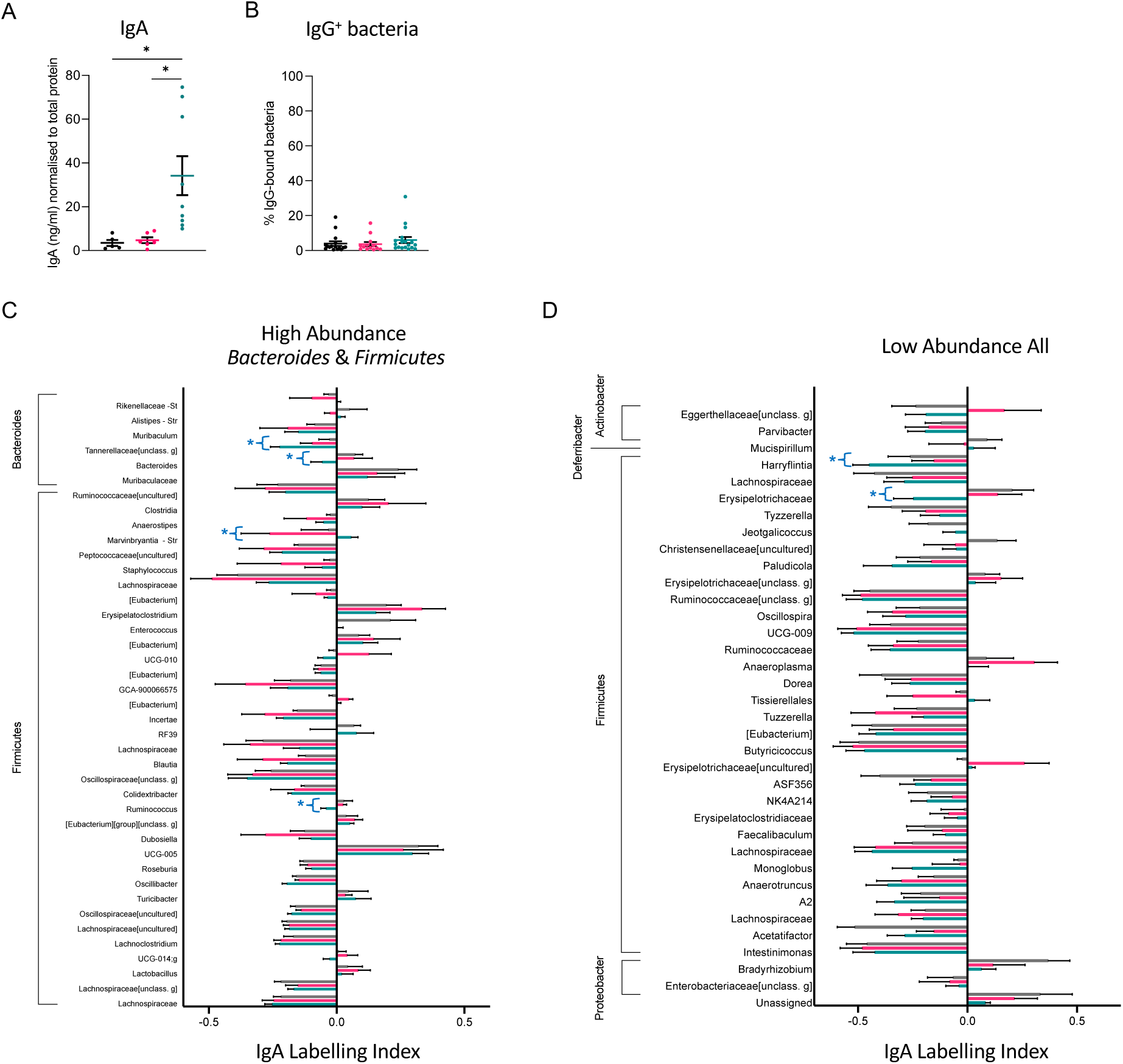
Mucosal antibody responses and binding to intestinal microbes in stroke. Related to Figure 4. A) Concentration of IgA normalized to total protein content of faecal pellet. B) IgG binding of fecal bacteria by flow cytometry. (C-D) IgA-binding index (Kau index) of genera of C) High abundance (rel. abundance >0.001) *Bacteroides* and *Firmicutes* genera and D) other less abundant (<0.001) genera detected. Data pooled from 2 independent experiment, n=7-9 per group. Data presented as mean +/-SEM. Statistical tests; A – one-way ANOVA w/ Tukey post-hoc, C-D Kruskal-Wallis test, * p <0.05 (A-B) or (C-D) * indicates difference within individual genera reached statistical significance.

**Figure S5.**
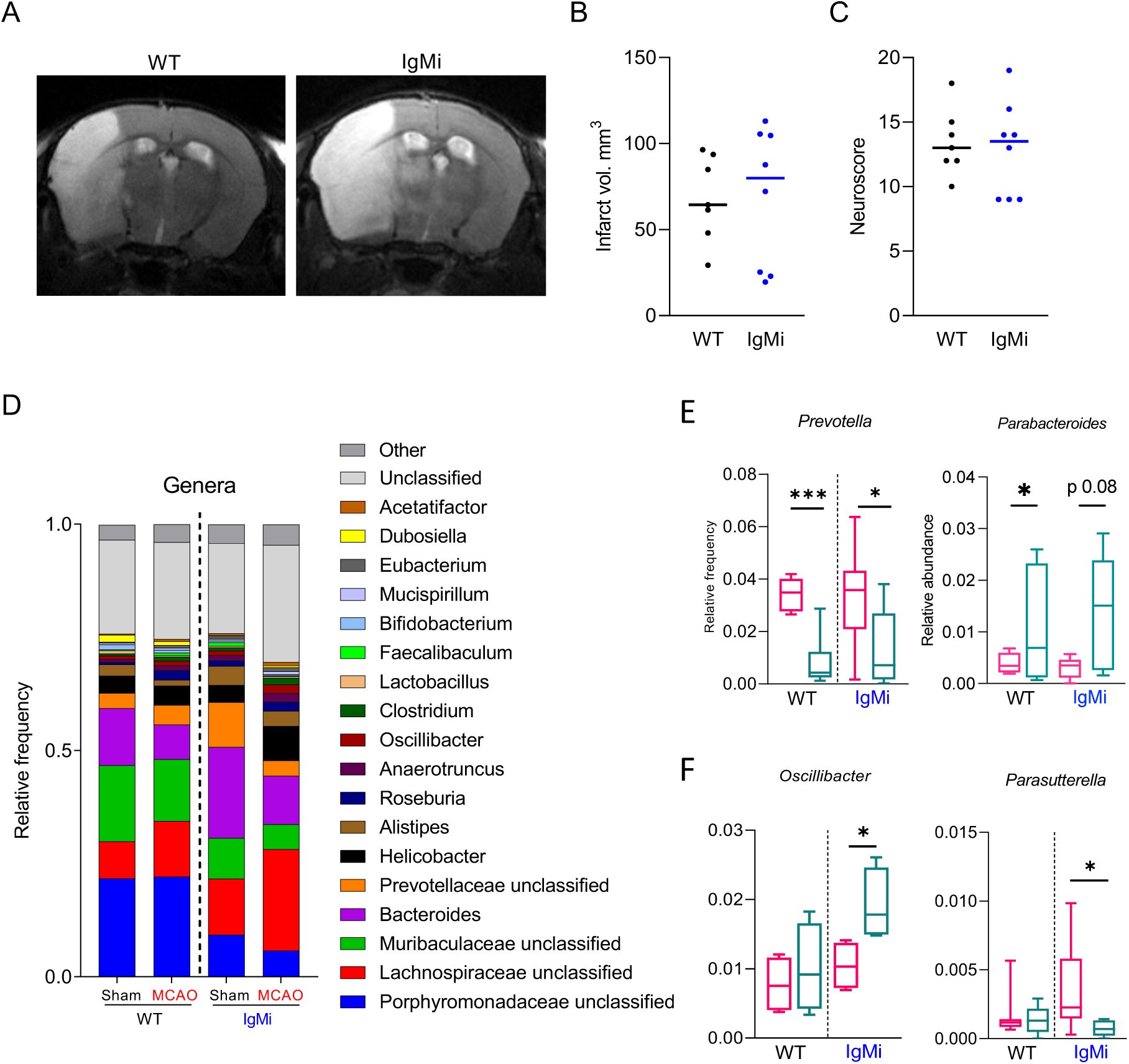
Stroke severity and microbial changes in IgMi mice. Related to Figure 5. A) Representative MRI scans of the brain of WT and IgMi mice 48h following MCAO. B) infarct volume and C) Neuroscore from WT and IgMi mice following stroke. (D-F) shotgun metagenomic sequencing of fecal microbiota isolated from IgMi and WT mice 48h after undergoing sham or MCAO surgery. D) Stacked bar charts representing average composition of microbiota at Genera level in WT and IgMi mice undergoing sham or MCAO. (E-F) Select individual genera from sham and stroke WT and IgMi mice, data pooled from 2 independent experiments, n=7-8. Data presented as mean +/-SEM, * p <0.05, ** p <0.01. Statistical tests; E-F – one-way ANOVA w/ Tukey post-hoc.

## References

1. Ashburner, M., C.A. Ball, J.A. Blake, D. Botstein, H. Butler, J.M. Cherry, A.P. Davis, K. Dolinski, S.S. Dwight, J.T. Eppig, M.A. Harris, D.P. Hill, L. Issel-Tarver, A. Kasarskis, S. Lewis, J.C. Matese, J.E. Richardson, M. Ringwald, G.M. Rubin, and G. Sherlock. 2000. Gene ontology: tool for the unification of biology. The Gene Ontology Consortium. Nat Genet 25:25–29.

2. Association, S. 2018. State of the nation: Stroke Statistics. In.

3. Belkaid, Y., and T.W. Hand. 2014. Role of the microbiota in immunity and inflammation. Cell 157:121–141.

4. Benakis, C., D. Brea, S. Caballero, G. Faraco, J. Moore, M. Murphy, G. Sita, G. Racchumi, L. Ling, E.G. Pamer, C. Iadecola, and J. Anrather. 2016. Commensal microbiota affects ischemic stroke outcome by regulating intestinal gammadelta T cells. Nat Med 22:516–523.

5. Benakis, C., and A. Liesz. 2022. The gut-brain axis in ischemic stroke: its relevance in pathology and as a therapeutic target. Neurol Res Pract 4:57.

6. Benakis, C., C. Martin-Gallausiaux, J.P. Trezzi, P. Melton, A. Liesz, and P. Wilmes. 2020a. The microbiome-gut-brain axis in acute and chronic brain diseases. Curr Opin Neurobiol 61:1–9.

7. Benakis, C., C. Poon, D. Lane, D. Brea, G. Sita, J. Moore, M. Murphy, G. Racchumi, C. Iadecola, and J. Anrather. 2020b. Distinct Commensal Bacterial Signature in the Gut Is Associated With Acute and Long-Term Protection From Ischemic Stroke. Stroke 51:1844–1854.

8. Brea, D., C. Poon, C. Benakis, G. Lubitz, M. Murphy, C. Iadecola, and J. Anrather. 2021. Stroke affects intestinal immune cell trafficking to the central nervous system. Brain Behav Immun 96:295–302.

9. Brichacek, A.L., D.C. Nwafor, S.A. Benkovic, S. Chakraborty, S.M. Kenney, M.E. Mace, S. Jun, C.A. Gambill, W. Wang, H. Hu, X. Ren, J.M. Povroznik, E.B. Engler-Chiurazzi, D.A. Primerano, J. Denvir, R. Percifield, A. Infante, J. Franko, R. Schafer, D.E. Gemoets, and C.M. Brown. 2020. Experimental Stroke Induces Chronic Gut Dysbiosis and Neuroinflammation in Male Mice. bioRxiv 2020.2004.2029.069575.

10. Brioschi, S., W.L. Wang, V. Peng, M. Wang, I. Shchukina, Z.J. Greenberg, J.K. Bando, N. Jaeger, R.S. Czepielewski, A. Swain, D.A. Mogilenko, W.L. Beatty, P. Bayguinov, J.A.J. Fitzpatrick, L.G. Schuettpelz, C.C. Fronick, I. Smirnov, J. Kipnis, V.S. Shapiro, G.F. Wu, S. Gilfillan, M. Cella, M.N. Artyomov, S.H. Kleinstein, and M. Colonna. 2021. Heterogeneity of meningeal B cells reveals a lymphopoietic niche at the CNS borders. Science 373:

11. Bunker, J.J., and A. Bendelac. 2018. IgA Responses to Microbiota. Immunity 49:211–224.

12. Camara-Lemarroy, C.R., B.E. Ibarra-Yruegas, and F. Gongora-Rivera. 2014. Gastrointestinal complications after ischemic stroke. J Neurol Sci 346:20–25.

13. Catanzaro, J.R., J.D. Strauss, A. Bielecka, A.F. Porto, F.M. Lobo, A. Urban, W.B. Schofield, and N.W. Palm. 2019. IgA-deficient humans exhibit gut microbiota dysbiosis despite secretion of compensatory IgM. Sci Rep 9:13574.

14. Clark, W., L. Gunion-Rinker, N. Lessov, and K. Hazel. 1998. Citicoline treatment for experimental intracerebral hemorrhage in mice. Stroke 29:2136–2140.

15. Clatworthy, M.R. 2021. The meninges-a cradle and school for nurturing and educating developing B cells. Immunity 54:2688–2690.

16. Consortium, G.O. 2021. The Gene Ontology resource: enriching a GOld mine. Nucleic Acids Res 49:D325–D334.

17. Crapser, J., R. Ritzel, R. Verma, V.R. Venna, F. Liu, A. Chauhan, E. Koellhoffer, A. Patel, A. Ricker, K. Maas, J. Graf, and L.D. McCullough. 2016. Ischemic stroke induces gut permeability and enhances bacterial translocation leading to sepsis in aged mice. Aging (Albany NY) 8:1049–1063.

18. Cui, W., L. Xu, L. Huang, Y. Tian, Y. Yang, Y. Li, and Q. Yu. 2023. Changes of gut microbiota in patients at different phases of stroke. CNS Neurosci Ther 29:3416–3429.

19. Delgado Jimenez R., and C. Benakis. 2021. The Gut Ecosystem: A Critical Player in Stroke. Neuromolecular Med 23:236–241.

20. Diaz-Marugan, L., M. Gallizioli, L. Marquez-Kisinousky, S. Arboleya, A. Mastrangelo, F. Ruiz-Jaen, J. Pedragosa, C. Casals, F.J. Morales, S. Ramos-Romero, S. Traserra, C. Justicia, M. Gueimonde, M. Jimenez, J.L. Torres, X. Urra, A. Chamorro, D. Sancho, C.G. de Los Reyes-Gavilan, F. Miro-Mur, and A.M. Planas. 2023. Poststroke Lung Infection by Opportunistic Commensal Bacteria Is Not Mediated by Their Expansion in the Gut Microbiota. Stroke 54:1875–1887.

21. Doyle, K.P., L.N. Quach, M. Sole, R.C. Axtell, T.V. Nguyen, G.J. Soler-Llavina, S. Jurado, J. Han, L. Steinman, F.M. Longo, J.A. Schneider, R.C. Malenka, and M.S. Buckwalter. 2015. B-lymphocyte-mediated delayed cognitive impairment following stroke. J Neurosci 35:2133–2145.

22. Durgan, D.J., J. Lee, L.D. McCullough, and R.M. Bryan, Jr. 2019. Examining the Role of the Microbiota-Gut-Brain Axis in Stroke. Stroke 50:2270–2277.

23. Feigin, V.L., M. Brainin, B. Norrving, S. Martins, R.L. Sacco, W. Hacke, M. Fisher, J. Pandian, and P. Lindsay. 2022. World Stroke Organization (WSO): Global Stroke Fact Sheet 2022. Int J Stroke 17:18–29.

24. Feigin, V.L., M.H. Forouzanfar, R. Krishnamurthi, G.A. Mensah, M. Connor, D.A. Bennett, A.E. Moran, R.L. Sacco, L. Anderson, T. Truelsen, M. O’Donnell, N. Venketasubramanian, S. Barker-Collo, C.M. Lawes, W. Wang, Y. Shinohara, E. Witt, M. Ezzati, M. Naghavi, C. Murray, I. Global Burden of Diseases, S. Risk Factors, and G.B.D.S.E.G. the. 2014. Global and regional burden of stroke during 1990-2010: findings from the Global Burden of Disease Study 2010. *Lancet* 383:245-254.

25. Fitzpatrick, Z., G. Frazer, A. Ferro, S. Clare, N. Bouladoux, J. Ferdinand, Z.K. Tuong, M.L. Negro-Demontel, N. Kumar, O. Suchanek, T. Tajsic, K. Harcourt, K. Scott, R. Bashford-Rogers, A. Helmy, D.S. Reich, Y. Belkaid, T.D. Lawley, D.B. McGavern, and M.R. Clatworthy. 2020. Gut-educated IgA plasma cells defend the meningeal venous sinuses. Nature 587:472–476.

26. Fitzpatrick, Z., N. Ghabdan Zanluqui, J.S. Rosenblum, Z.K. Tuong, C.Y.C. Lee, V. Chandrashekhar, M.L. Negro-Demontel, A.P. Stewart, D.A. Posner, M. Buckley, K.S.J. Allinson, P. Mastorakos, P. Chittiboina, D. Maric, D. Donahue, A. Helmy, T. Tajsic, J.R. Ferdinand, A. Portet, A. Penalver, E. Gillman, Z. Zhuang, M.R. Clatworthy, and D.B. McGavern. 2024. Venous-plexus-associated lymphoid hubs support meningeal humoral immunity. Nature 628:612–619.

27. Ge, Y., M. Zadeh, C. Yang, E. Candelario-Jalil, and M. Mohamadzadeh. 2022. Ischemic Stroke Impacts the Gut Microbiome, Ileal Epithelial and Immune Homeostasis. iScience 25:105437.

28. Houlden, A., M. Goldrick, D. Brough, E.S. Vizi, N. Lenart, B. Martinecz, I.S. Roberts, and A. Denes. 2016. Brain injury induces specific changes in the caecal microbiota of mice via altered autonomic activity and mucoprotein production. Brain Behav Immun 57:10–20.

29. Huus, K.E., C. Petersen, and B.B. Finlay. 2021. Diversity and dynamism of IgA-microbiota interactions. Nat Rev Immunol 21:514–525.

30. Iadecola, C., and J. Anrather. 2011. The immunology of stroke: from mechanisms to translation. Nat Med 17:796–808.

31. Iadecola, C., M.S. Buckwalter, and J. Anrather. 2020. Immune responses to stroke: mechanisms, modulation, and therapeutic potential. J Clin Invest 130:2777–2788.

32. Kau, A.L., J.D. Planer, J. Liu, S. Rao, T. Yatsunenko, I. Trehan, M.J. Manary, T.C. Liu, T.S. Stappenbeck, K.M. Maleta, P. Ashorn, K.G. Dewey, E.R. Houpt, C.S. Hsieh, and J.I. Gordon. 2015. Functional characterization of IgA-targeted bacterial taxa from undernourished Malawian children that produce diet-dependent enteropathy. Sci Transl Med 7:276ra224.

33. Lin, C.J., J.W. Hung, C.Y. Cho, C.Y. Tseng, H.Y. Chen, F.C. Lin, and C.Y. Li. 2013. Poststroke constipation in the rehabilitation ward: incidence, clinical course and associated factors. Singapore Med J 54:624–629.

34. Liu, C., X. Cheng, S. Zhong, Z. Liu, F. Liu, X. Lin, Y. Zhao, M. Guan, T. Xiao, J. Jolkkonen, Y. Wang, and C. Zhao. 2022. Long-term modification of gut microbiota by broad-spectrum antibiotics improves stroke outcome in rats. Stroke Vasc Neurol 7:381–389.

35. Longa, E.Z., P.R. Weinstein, S. Carlson, and R. Cummins. 1989. Reversible middle cerebral artery occlusion without craniectomy in rats. Stroke 20:84–91.

36. Luck, H., S. Khan, J.H. Kim, J.K. Copeland, X.S. Revelo, S. Tsai, M. Chakraborty, K. Cheng, Y. Tao Chan, M.K. Nohr, X. Clemente-Casares, M.C. Perry, M. Ghazarian, H. Lei, Y.H. Lin, B. Coburn, A. Okrainec, T. Jackson, S. Poutanen, H. Gaisano, J.P. Allard, D.S. Guttman, M.E. Conner, S. Winer, and D.A. Winer. 2019. Gut-associated IgA(+) immune cells regulate obesity-related insulin resistance. Nat Commun 10:3650.

37. Lycke, N.Y., and M. Bemark. 2012. The role of Peyer’s patches in synchronizing gut IgA responses. Front Immunol 3:329.

38. Lynch, S.V., and O. Pedersen. 2016. The Human Intestinal Microbiome in Health and Disease. N Engl J Med 375:2369–2379.

39. Macpherson, A.J., K.D. McCoy, F.E. Johansen, and P. Brandtzaeg. 2008. The immune geography of IgA induction and function. Mucosal Immunol 1:11–22.

40. McCulloch, L., C.J. Smith, and B.W. McColl. 2017. Adrenergic-mediated loss of splenic marginal zone B cells contributes to infection susceptibility after stroke. Nat Commun 8:15051.

41. Nouraee, C., M. Fisher, M. Di Napoli, P. Salazar, T.D. Farr, A. Jafarli, and A.A. Divani. 2019. A Brief Review of Edema-Adjusted Infarct Volume Measurement Techniques for Rodent Focal Cerebral Ischemia Models with Practical Recommendations. J Vasc Interv Neurol 10:38–45.

42. Ortega, S.B., V.O. Torres, S.E. Latchney, C.W. Whoolery, I.Z. Noorbhai, K. Poinsatte, U.M. Selvaraj, M.A. Benson, A.J.M. Meeuwissen, E.J. Plautz, X. Kong, D.M. Ramirez, A.D. Ajay, J.P. Meeks, M.P. Goldberg, N.L. Monson, A.J. Eisch, and A.M. Stowe. 2020. B cells migrate into remote brain areas and support neurogenesis and functional recovery after focal stroke in mice. Proc Natl Acad Sci U S A 117:4983–4993.

43. Oyama, N., K. Winek, P. Backer-Koduah, T. Zhang, C. Dames, M. Werich, O. Kershaw, C. Meisel, A. Meisel, and U. Dirnagl. 2018. Exploratory Investigation of Intestinal Function and Bacterial Translocation After Focal Cerebral Ischemia in the Mouse. Front Neurol 9:937.

44. Pabst, O. 2012. New concepts in the generation and functions of IgA. Nat Rev Immunol 12:821–832. Pabst, O., and E. Slack. 2020. IgA and the intestinal microbiota: the importance of being specific. *Mucosal Immunol* 13:12-21.

45. Penny, H.A., R.G. Domingues, M.Z. Krauss, F. Melo-Gonzalez, M.A.E. Lawson, S. Dickson, J. Parkinson, M. Hurry, C. Purse, E. Jegham, C. Godinho-Silva, M. Rendas, H. Veiga-Fernandes, D.A. Bechtold, R.K. Grencis, K.M. Toellner, A. Waisman, J.R. Swann, J.E. Gibbs, and M.R. Hepworth. 2022. Rhythmicity of intestinal IgA responses confers oscillatory commensal microbiota mutualism. Sci Immunol 7:eabk2541.

46. Percie du Sert, N., A. Ahluwalia, S. Alam, M.T. Avey, M. Baker, W.J. Browne, A. Clark, I.C. Cuthill, U. Dirnagl, M. Emerson, P. Garner, S.T. Holgate, D.W. Howells, V. Hurst, N.A. Karp, S.E. Lazic, K. Lidster, C.J. MacCallum, M. Macleod, E.J. Pearl, O.H. Petersen, F. Rawle, P. Reynolds, K. Rooney, E.S. Sena, S.D. Silberberg, T. Steckler, and H. Würbel. 2020. Reporting animal research: Explanation and elaboration for the ARRIVE guidelines 2.0. PLoS Biol 18:e3000411.

47. Probstel, A.K., X. Zhou, R. Baumann, S. Wischnewski, M. Kutza, O.L. Rojas, K. Sellrie, A. Bischof, K. Kim, A. Ramesh, R. Dandekar, A.L. Greenfield, R.D. Schubert, J.E. Bisanz, S. Vistnes, K. Khaleghi, J. Landefeld, G. Kirkish, F. Liesche-Starnecker, V. Ramaglia, S. Singh, E.B. Tran, P. Barba, K. Zorn, J. Oechtering, K. Forsberg, L.R. Shiow, R.G. Henry, J. Graves, B.A.C. Cree, S.L. Hauser, J. Kuhle, J.M. Gelfand, P.M. Andersen, J. Schlegel, P.J. Turnbaugh, P.H. Seeberger, J.L. Gommerman, M.R. Wilson, L. Schirmer, and S.E. Baranzini. 2020. Gut microbiota-specific IgA(+) B cells traffic to the CNS in active multiple sclerosis. Sci Immunol 5:

48. Rigoni, R., E. Fontana, S. Guglielmetti, B. Fosso, A.M. D’Erchia, V. Maina, V. Taverniti, M.C. Castiello, S. Mantero, G. Pacchiana, S. Musio, R. Pedotti, C. Selmi, J.R. Mora, G. Pesole, P. Vezzoni, P.L. Poliani, F. Grassi, A. Villa, and B. Cassani. 2016. Intestinal microbiota sustains inflammation and autoimmunity induced by hypomorphic RAG defects. J Exp Med 213:355–375.

49. Rojas, O.L., A.K. Probstel, E.A. Porfilio, A.A. Wang, M. Charabati, T. Sun, D.S.W. Lee, G. Galicia, V. Ramaglia, L.A. Ward, L.Y.T. Leung, G. Najafi, K. Khaleghi, B. Garcillan, A. Li, R. Besla, I. Naouar, E.Y. Cao, P. Chiaranunt, K. Burrows, H.G. Robinson, J.R. Allanach, J. Yam, H. Luck, D.J. Campbell, D. Allman, D.G. Brooks, M. Tomura, R. Baumann, S.S. Zamvil, A. Bar-Or, M.S. Horwitz, D.A. Winer, A. Mortha, F. Mackay, A. Prat, L.C. Osborne, C. Robbins, S.E. Baranzini, and J.L. Gommerman. 2019. Recirculating Intestinal IgA-Producing Cells Regulate Neuroinflammation via IL-10. Cell 176:610–624 e618.

50. Rollenske, T., S. Burkhalter, L. Muerner, S. von Gunten, J. Lukasiewicz, H. Wardemann, and A.J. Macpherson. 2021. Parallelism of intestinal secretory IgA shapes functional microbial fitness. Nature 598:657–661.

51. Sahputra, R., J.C. Yam-Puc, A. Waisman, W. Muller, and K.J. Else. 2018. Evaluating the IgMi mouse as a novel tool to study B-cell biology. Eur J Immunol 48:2068–2071.

52. Schaller, B.J., R. Graf, and A.H. Jacobs. 2006. Pathophysiological changes of the gastrointestinal tract in ischemic stroke. Am J Gastroenterol 101:1655–1665.

53. Schulte-Herbruggen, O., D. Quarcoo, A. Meisel, and C. Meisel. 2009. Differential affection of intestinal immune cell populations after cerebral ischemia in mice. Neuroimmunomodulation 16:213–218.

54. Singh, V., S. Roth, G. Llovera, R. Sadler, D. Garzetti, B. Stecher, M. Dichgans, and A. Liesz. 2016. Microbiota Dysbiosis Controls the Neuroinflammatory Response after Stroke. J Neurosci 36:7428–7440.

55. Stanley, D., L.J. Mason, K.E. Mackin, Y.N. Srikhanta, D. Lyras, M.D. Prakash, K. Nurgali, A. Venegas, M.D. Hill, R.J. Moore, and C.H. Wong. 2016. Translocation and dissemination of commensal bacteria in post-stroke infection. Nat Med 22:1277–1284.

56. Stanley, D., R.J. Moore, and C.H.Y. Wong. 2018. An insight into intestinal mucosal microbiota disruption after stroke. Sci Rep 8:568.

57. Sun, H., M. Gu, Z. Li, X. Chen, and J. Zhou. 2021. Gut Microbiota Dysbiosis in Acute Ischemic Stroke Associated With 3-Month Unfavorable Outcome. Front Neurol 12:799222.

58. Suzuki, K., B. Meek, Y. Doi, M. Muramatsu, T. Chiba, T. Honjo, and S. Fagarasan. 2004. Aberrant expansion of segmented filamentous bacteria in IgA-deficient gut. Proc Natl Acad Sci U S A 101:1981–1986.

59. Tuz, A.A., A. Hasenberg, D.M. Hermann, M. Gunzer, and V. Singh. 2022. Ischemic stroke and concomitant gastrointestinal complications-a fatal combination for patient recovery. Front Immunol 13:1037330.

60. Waisman, A., M. Kraus, J. Seagal, S. Ghosh, D. Melamed, J. Song, Y. Sasaki, S. Classen, C. Lutz, F. Brombacher, L. Nitschke, and K. Rajewsky. 2007. IgG1 B cell receptor signaling is inhibited by CD22 and promotes the development of B cells whose survival is less dependent on Ig alpha/beta. J Exp Med 204:747–758.

61. Westendorp, W.F., P.J. Nederkoorn, J.D. Vermeij, M.G. Dijkgraaf, and D. van de Beek. 2011. Post-stroke infection: a systematic review and meta-analysis. BMC Neurol 11:110.

62. Xia, G.H., C. You, X.X. Gao, X.L. Zeng, J.J. Zhu, K.Y. Xu, C.H. Tan, R.T. Xu, Q.H. Wu, H.W. Zhou, Y. He, and J. Yin. 2019. Stroke Dysbiosis Index (SDI) in Gut Microbiome Are Associated With Brain Injury and Prognosis of Stroke. Front Neurol 10:397.

63. Xu, R., C. Tan, J. Zhu, X. Zeng, X. Gao, Q. Wu, Q. Chen, H. Wang, H. Zhou, Y. He, S. Pan, and J. Yin. 2019. Dysbiosis of the intestinal microbiota in neurocritically ill patients and the risk for death. Crit Care 23:195.

64. Yamashiro, K., R. Tanaka, T. Urabe, Y. Ueno, Y. Yamashiro, K. Nomoto, T. Takahashi, H. Tsuji, T. Asahara, and N. Hattori. 2017. Gut dysbiosis is associated with metabolism and systemic inflammation in patients with ischemic stroke. PLoS One 12:e0171521.

65. Yin, J., S.X. Liao, Y. He, S. Wang, G.H. Xia, F.T. Liu, J.J. Zhu, C. You, Q. Chen, L. Zhou, S.Y. Pan, and H.W. Zhou. 2015. Dysbiosis of Gut Microbiota With Reduced Trimethylamine-N-Oxide Level in Patients With Large-Artery Atherosclerotic Stroke or Transient Ischemic Attack. J Am Heart Assoc 4:

